# The molecular basis of integrated stress response silencing

**DOI:** 10.1101/2024.12.02.626349

**Authors:** Zhi Yang, Diane L. Haakonsen, Michael Heider, Samuel R. Witus, Alex Zelter, Tobias Beschauner, Michael J. MacCoss, Michael Rapé

## Abstract

Chronic stress response activation impairs cell survival and causes devastating degenerative diseases. To counteract this, cells deploy dedicated silencing factors, such as the E3 ligase SIFI that terminates the mitochondrial stress response. How a single enzyme can sense stress across cells and elicit timely stress response inactivation is poorly understood. Here, we report the structure of human SIFI, which revealed how this 1.3MDa complex can target hundreds of proteins for accurate stress response silencing. SIFI attaches the first ubiquitin to substrates using flexible domains within an easily accessible scaffold, yet builds linkage-specific ubiquitin chains at distinct, sterically restricted elongation modules in its periphery. Ubiquitin handover via a ubiquitin-like domain couples versatile substrate modification to precise chain elongation. Stress response silencing therefore exploits a catalytic mechanism that is geared to process many diverse proteins and hence allows a single enzyme to monitor and, if appropriate, modulate a complex cellular state.

## Introduction

The resilience of metazoan development relies on stress response pathways that detect and mitigate mutational, physiological, or environmental challenges onto cellular processes ^1–3^. Providing a key example, mitochondrial import stress activates the HRI kinase, which phosphorylates the translation initiation factor eIF2α to stall the synthesis of mitochondrial proteins ^4–6^. In this manner, HRI reduces the cytoplasmic load of aggregation-prone proteins and provides cells with time to restore mitochondrial integrity. HRI is one of four kinases of the integrated stress response that allows cells to overcome a wide range of deleterious conditions ^1,7–9^.

As stress responses put core processes on hold, their prolonged activation can trigger cell death and tissue degeneration ^8^. To prevent this from happening, cells turn off stress responses using dedicated silencing factors ^10^. Upon restoration of mitochondrial import, the E3 ubiquitin ligase SIFI (***si***lencing ***f***actor of the ***i***ntegrated stress response) marks HRI and its activator DELE1 for proteasomal degradation to terminate eIF2α phosphorylation and restart protein synthesis ^10^. Deletion of the SIFI subunit UBR4 disrupts heart, brain and yolk-sac development, thereby causing embryonic lethality ^11,12^, and mutations in *UBR4* lead to ataxia and early-onset dementia ^13–15^. As genetic or pharmacological inactivation of HRI rescued *UBR4*-deficient cells ^10^, stress response silencing is critical for cell and tissue homeostasis.

To ensure timely silencing, SIFI must act across cells to sense if stress is still present. SIFI accomplishes this task by recognizing mitochondrial precursors that only accumulate in the cytoplasm during stress ^10,16–18^, and it also detects cleaved proteins that can be released from stressed mitochondria ^4,5,10,19^. These targets compete with HRI for access to SIFI and thus delay stress response silencing until stress has been resolved ^10^. How SIFI can process hundreds of proteins that differ in size from ∼50 to ∼2500 residues and adopt a wide range of conformations is un-known. More generally, how a single E3 ligase can evaluate a global cellular state remains to be elucidated.

Here, we report the structure of SIFI and reveal the molecular basis of stress response silencing. SIFI attaches the first ubiquitin to substrates using flexible domains within an open scaffold, while it extends K48-linked polymers at separate modules built around a sterically restricted E2. SIFI’s catalytic centers communicate via ubiquitin handover orchestrated by a ubiquitin-like domain. Silencing of the integrated stress response therefore relies on a catalytic mechanism that couples versatile substrate modification to linkage-specific chain elongation, which allows a single enzyme to process many diverse proteins for timely stress response silencing.

## Results

### Structure of endogenous SIFI

To determine the structure of SIFI, we purified the endogenous complex by immunoprecipitating its largest subunit, UBR4, followed by size exclusion chromatography (**Extended Data** Fig. 1a). Having ensured that SIFI was active (**Extended Data** Fig. 1b), we used single particle cryo-EM to resolve a partial map of UBR4 and associated proteins to an overall 3.1Å resolution (**Fig. 1a**; **Extended Data** Fig. 1c). Purification of SIFI via its subunit KCMF1 led to a complementary map of the N-terminal half of UBR4 at an overall 3.4Å resolution (**Fig. 1a**; **Extended Data** Fig. 1d). The C-terminal region of UBR4, which includes its hemi-RING ^20^, was not resolved until one of SIFI’s E2 enzymes, UBE2A, was supplemented prior to cryo-EM analysis (**Extended Data** Fig. 1e). Lastly, we integrated AlphaFold models of specific domains into less well-resolved regions and combined all maps to build a near-complete structure of human SIFI (**Fig. 1b, c**; **Supplementary Movie 1**).

**Figure 1:**
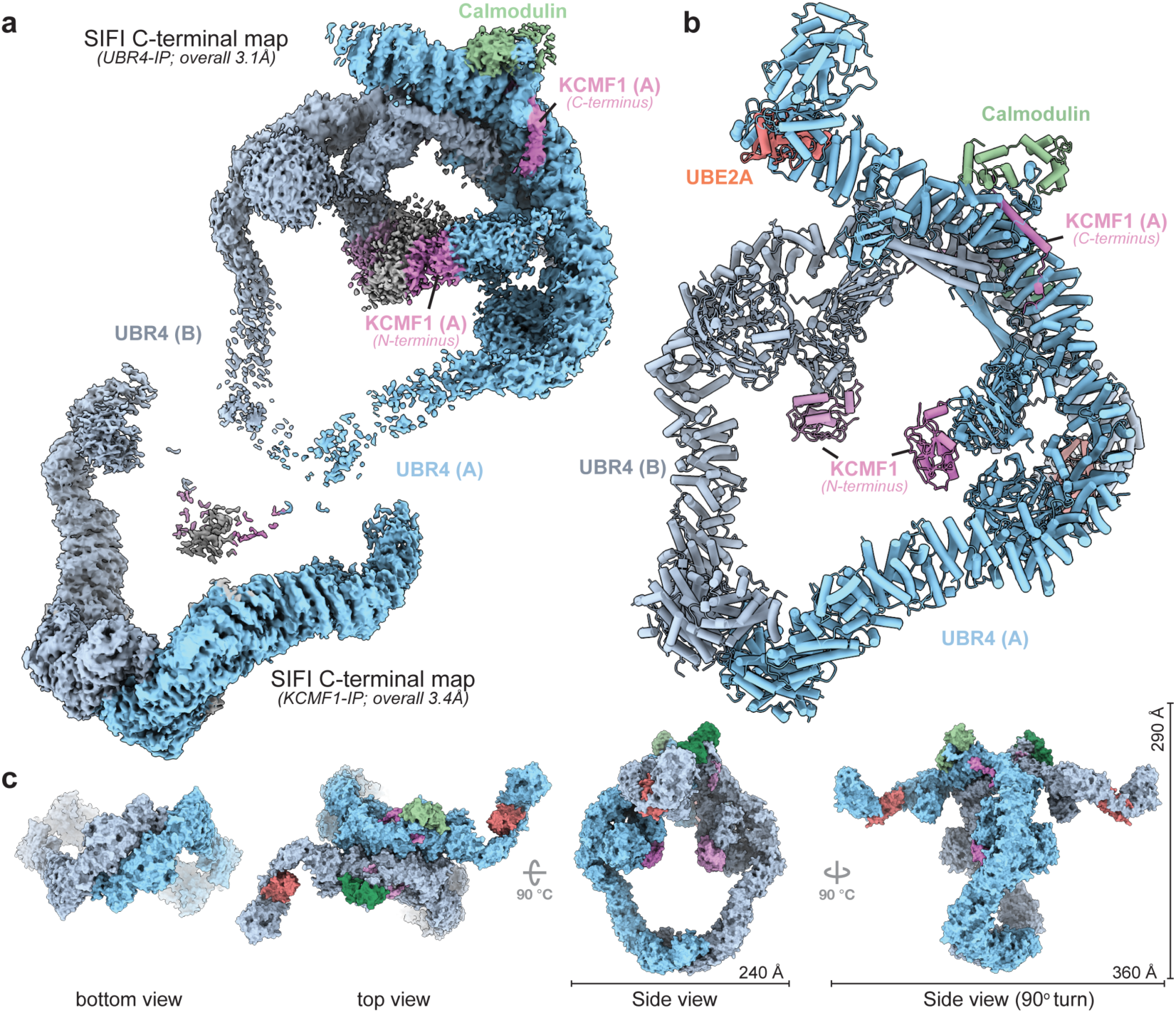
Cryo-EM structure of human SIFI. **a.** Cryo-EM maps of SIFI highlighting its C-terminal partial map (top left, endogenous SIFI complex) and N-terminal partial map (immunoprecipitation of KCMF1, bottom left). **b.** Composite structure of human SIFI built from partial maps, including two subunits each of UBR4 (blue), KCMF1 (purple), calmodulin (green), and UBE2A (red). **c.** Surface representations of the SIFI complex viewed from the N- and C-terminal dimerization regions, as well as views from above and the side of ring’s plane. All components are color-coded as in panels **a** and **b**.

Our structure showed that SIFI forms an antiparallel dimer of heterotrimers that comprises two copies each of the E3 ligase UBR4, the E3 ligase KCMF1, and the calcium-binding chaperone calmodulin (**Fig. 1b, c**; **Extended Data** Fig. 1f). UBE2A was captured by the C-terminal region of each UBR4 subunit that became ordered in its presence. Together, these proteins assemble into a ∼1.3MDa structure that is built around an open twisted-ring scaffold (292Åx229Å) and two arms that extend to the side and span ∼356Å. SIFI compares in size to 80S ribosomes and 26S proteasomes (**Extended Data** Fig. 1g).

SIFI’s scaffold is primarily composed of α-helical armadillo repeats of UBR4 and relies on antiparallel dimer interfaces at its N- and C-terminal regions with buried surface areas of ∼2,160Å^2^ and ∼4,212Å^2^, respectively (**Fig. 1b, c**). The N-terminal dimer interface is mediated by a domain-swapped helix from one protomer that bundles with three helices of the second, as well as two Arg residues that interlock the dimer via polar interactions (**Extended Data** Fig. 2a). The C-terminal interface is centered on extended interactions between helical repeats from each protomer that are cemented by horizontal coiled-coils at the inner side of the scaffold (**Extended Data Fig. 2b**). Revealing flexibility of the scaffold, the dimerization interfaces show ∼10° rotations towards each other, which occur around a hinge close to UBR4-A2581 that is mutated in ataxia and R2584 that is altered in cancer ^21,22^ (**Extended Data** Fig. 2c; **Supplementary Movie 2**).

Calmodulin docks onto the outer rim of the scaffold, close to a hinge around UBR4-G4301 that connects the C-terminal dimer interface to the catalytic module of UBR4 (**Fig. 2a-c**). Calmodulin’s C-terminal four-helix lobe, which binds calcium, engages an exposed UBR4 helix that fits the calmodulin-binding consensus and contains Arg residues mutated in cancer cells and ataxia ^22–24^ (**Fig. 2a, b**). Hydrophobic calmodulin residues surround Trp4105 of UBR4 in a manner consistent with other calcium-activated structures ^24,25^. Calmodulin’s N-lobe, which is free of calcium, forms a bundle with two helices of the UBR4 scaffold (**Fig. 2b, c**). In this manner, calmodulin creates an intramolecular bridge that stabilizes a region of UBR4 that we refer to as ‘lid’ and that anchors KCMF1 to the scaffold, as described below. These findings explain why deletion of UBR4 residues that include its exposed helix prevented calmodulin integration into SIFI and stress response silencing ^10,26^.

**Figure 2:**
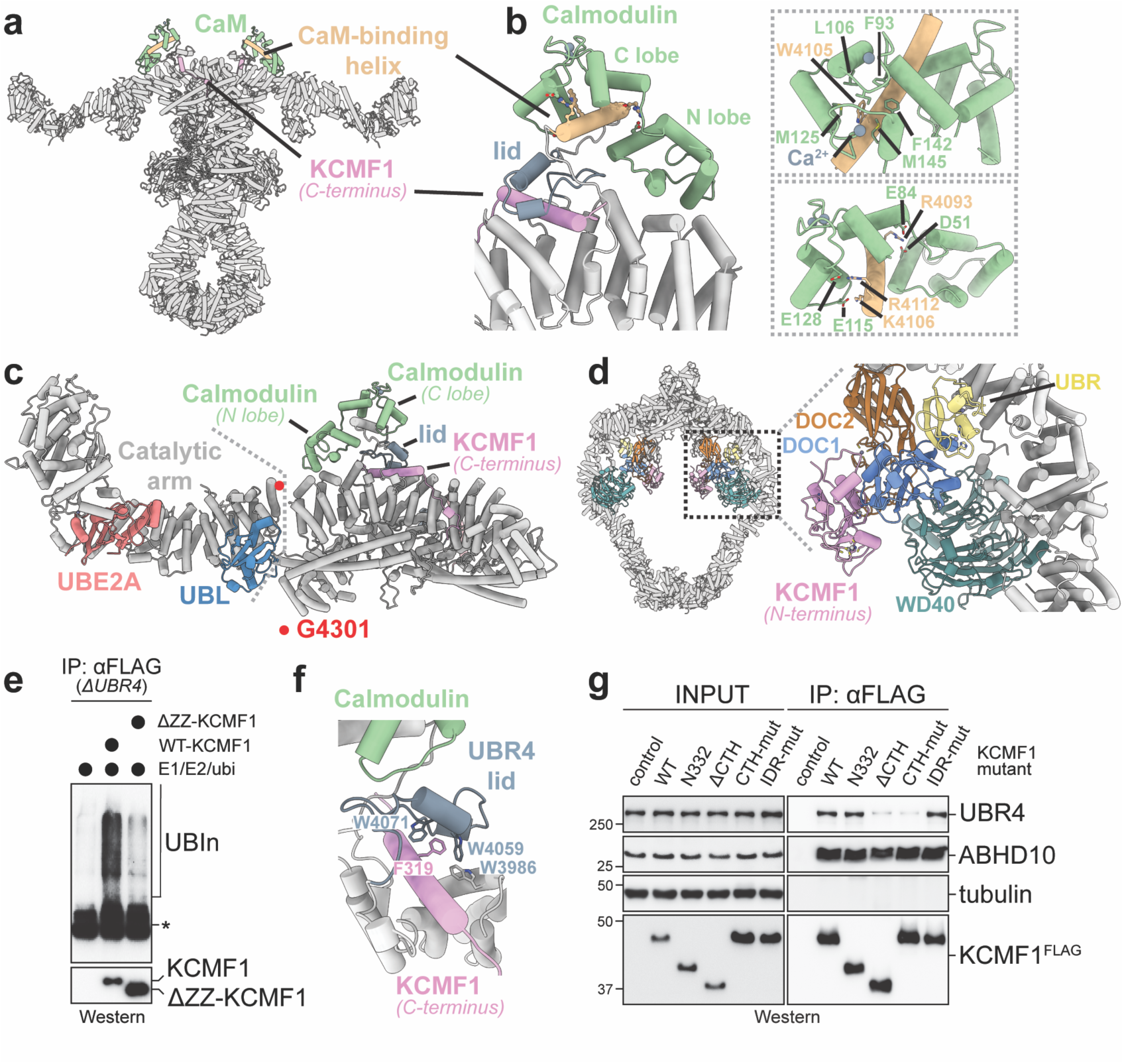
Structural arrangements of SIFI modules. **a.** Calmodulin (green) binds to the outer rim of the SIFI scaffold above the C-terminal dimerization interface of UBR4. **b.** The Ca^2+^-free N-terminal lobe of calmodulin (green) forms a helix bundle with UBR4 (light gray), while the Ca^2+^-bound C-terminal lobe of calmodulin engages a calmodulin-binding motif of UBR4 (gold), situated atop a UBR4 lid structure (blue gray) that encircles the C-terminal α-helix of KCMF1 (plum). Conserved hydrophobic and polar residues involved in the interaction are highlighted within dashed-line boxes (*right panels*). **c.** Calmodulin stabilizes the base of the UBR4 C-terminal catalytic arm, including hemi-RING, UBE2A (coral), and UBL (blue), allowing flexible movement of the arm around residue Gly4301 (red dot) of UBR4. **d.** Left: two copies each of the protein interaction modules (colored) of SIFI are located at the center of the ring architecture. Right: a close-up view of these modules, including the DOC2 domain (brown), WD40 domain (teal), and the KCMF1^N138^-DOC1-UBR subcomplex (plum, blue, and yellow, respectively). Within this subcomplex, the two subdomains of KCMF1^N138^, the ZZ-type and C2H2-type domains, are situated. **e.** The N-terminal ZZ domain of KCMF1 is required for ubiquitylation activity. WT- or ΔZZ-KCMF1 were affinity-purified from *ΔUBR4* cells to prevent UBR4-dependent ubiquitylation and incubated with E1, E2 (UBE2D3, UBE2A) and ubiquitin. Ubiquitylation activity was detected by formation of high-MW ubiquitin conjugates, using αUbiquitin-Western. Similar results in n=2 independent experiments. **f.** C-terminal helix of KCMF1 (plum) is anchored within the α-helical bundles of UBR4 Armadillo repeats (light gray) through the UBR4 lid (blue-gray). A conserved Phe319 of KCMF1 (plum), surrounded by three Trp residues of UBR4 (blue and light gray), is highlighted. **g.** Deletion or mutation (R316E/F319E/L323E/L325E) of the C-terminal helix impedes KCMF1 integration into SIFI, as shown by KCMF1^FLAG^ affinity-purification and detection of UBR4. A Q226/228/229/231/233E mutation in an unresolved linker of KCMF1 (IDR-mut.) did not affect integration into SIFI. ABHD10 binds to the ZZ-domain of KCMF1 and is used as a positive control. Similar results in n=2 independent experiments.

Tethered to the scaffold’s interior are a UBR-, WD40-, and two DOC-homologous β-sandwich domains of each UBR4 subunit (**Fig. 1b**; **Fig. 2d**). Their lower local resolution indicates that these modules are flexibly attached to the scaffold (**Extended Data** Fig. 2d). The WD40 repeats reside symmetrically on either side of the ring, where they are supported by a helix protruding from the scaffold. A second short helix is predicted to plug each β-propeller using loops that often recognize binding partners of WD40 repeats (**Extended Data** Fig. 2e). The DOC1 domains are positioned near each β-propeller and engage the UBR box of UBR4 and the N-terminal region of KCMF1 to form a three-component subcomplex, referred to as the KCMF1 module, that fits well into the local ∼16Å map (**Fig. 2d**). Indicative of its inherent flexibility, 3D classification analyses indicated that this subcomplex can adopt distinct positions on SIFI (**Supplementary Movie 1**). The DOC2 domains reside next to the KCMF1 module, where they combine a β-sandwich characteristic of DOC domains with two exposed Zn^2+^-coordinating loops that face the central cavity, forming another potential protein-binding interface (**Extended Data** Fig. 2f). Chemical cross-linking combined with mass spectrometry confirmed the position of these interaction modules within the center of SIFI (**Extended Data** Fig. 2g).

SIFI possesses two subunits, KCMF1 and UBR4, with unconventional E3 ligase motifs. In KCMF1, we found that the N-terminal ZZ-domain, which shares structural similarity to RING domains, is required for ubiquitylation (**Fig. 2e**). This domain as well as two predicted C2H2-type Zinc-fingers are near the center of the scaffold, where they bind UBR4’s DOC1 domain (**Fig. 2d**). Following an unresolved linker, KCMF1 is further anchored in SIFI above the C-terminal dimer interface of UBR4 through a helix that is enclosed by the UBR4 lid (**Fig. 2f**). The UBR4 lid engages the KCMF1 helix via extensive van der Waals and polar interactions, including a group of Trp residues that encircle the conserved Phe319 of KCMF1 (**Fig. 2f**; **Extended Data** Fig. 3a). Mutation of interface residues or deletion of N- or C-terminal regions of KCMF1 impaired its retention in SIFI (**Fig. 2g**; **Extended Data** Fig. 3b), while excision of the DOC1 domain in *UBR4* had previously been shown to disrupt KCMF1 binding ^10^.

The ubiquitylation modules of UBR4 are found within the SIFI arms that make a ∼90° turn to extend ∼166Å above the scaffold (**Fig. 1b, c**) ^20^. These catalytic sites include a 10-turn α-helix, a hemi-RING and a UZI domain in which several ataxia mutations are found ^15,22^. Based on X-ray structures and AlphaFold models ^20^, the long helix binds to the backside of the E2 and together with the hemi-RING captures UBE2A in an embrace that likely explains why this region was only visible when SIFI was incubated with its E2. Close to this module is a ubiquitin-like (UBL) domain that is tethered to the UBR4 scaffold through long linkers rich in Asp and Glu residues (**Fig. 2c**). As described below, the UBL domain plays an important role in ubiquitin chain formation.

Together, our structure reveals that SIFI contains several interaction motifs within a large and easily accessible scaffold. Two catalytic sites are provided by the KCMF1 subunits at the center of this structure, while two further ubiquitylation modules reside within the peripheral SIFI arms. With four ubiquitylation centers and a flexible architecture, SIFI appears to be optimally configured to process many substrates for efficient stress response silencing.

### Substrate engagement in the center of the SIFI scaffold

To prevent premature stress response silencing, SIFI must monitor stress across the entire cell. To this end, SIFI detects unimported or cleaved proteins that accumulate in the cytoplasm during stress ^10^, but how it can engage so many diverse substrates is unknown. In the cryo-EM map of endogenous SIFI, we noticed a low-resolution density that was attached to KCMF1 and could not be ascribed to any SIFI component (**Fig. 3a**). To identify this factor, we compared SIFI immunoprecipitates from WT and *ΔKCMF1* cells by mass spectrometry and found that two mitochondrial proteins, ABHD10 and the less abundant NIPSNAP3A, were lost in the absence of KCMF1 (**Extended Data Fig. 4a**). The same proteins were found to bind KCMF1 in *ΔUBR4* cells (**Extended Data** Fig. 4b). AlphaFold models showed that ABHD10 dimers fit well into the central density observed in the cryo-EM map (**Fig. 3a**). Intriguingly, ABHD10 was efficiently ubiquitylated by SIFI (**Extended Data** Fig. 4c), suggesting that it was a substrate, rather than subunit, of the E3 ligase.

**Figure 3:**
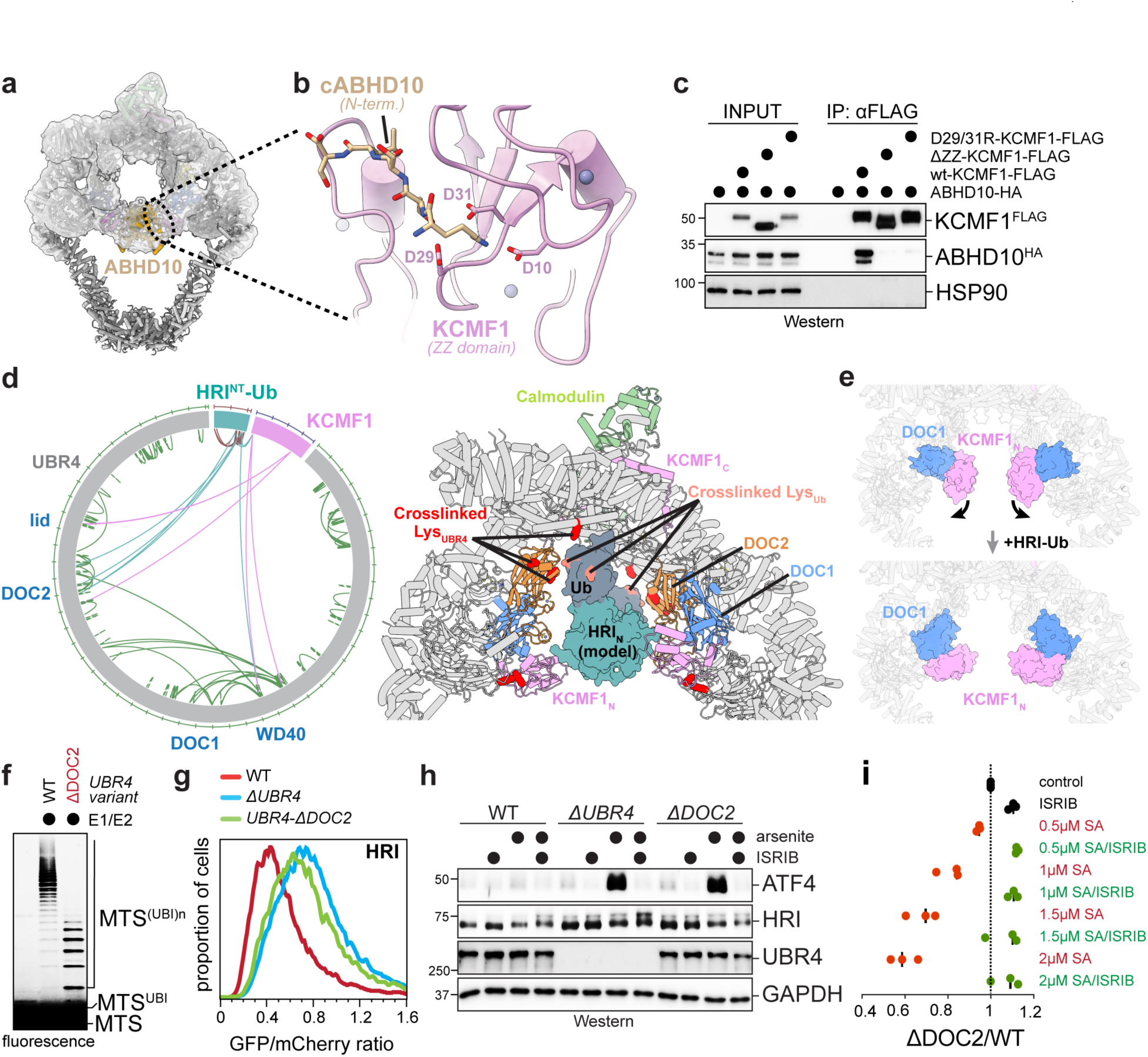
SIFI binds substrates at the center of its twisted-ring scaffold. **a.** Overview of SIFI highlighting a central low-resolution density ascribed to dimers of ABHD10. **b.** AF2 model of ABHD10’s cleaved N-terminus fitting into the N-degron binding pocket within the ZZ-domain of KCMF1. D29 and D31 of KCMF1, which interact with the first Lys of the ABHD10 N-degron, are highlighted. **c.** Deletion of KCMF1’s ZZ-domain or mutation of D29 and D31 ablates recognition of ABHD10. Immunoprecipitation of KCMF1^FLAG^ variants co-overexpressed with ABHD10^HA^ from *ΔUBR4* 293T cells. Experiment was performed once. **d.** *Left*: diagram of cross-linking mass spectrometry of the SIFI∼HRI-Ub com-plex, highlighting directly cross-linked residues from UBR4, KCMF1, and HRI∼Ub. HRI^NT^ contains scarce Lys residues, and peptides of HRI^NT^ were not detected in mass spectrometry. *Right*: a model of dimerized HRI∼Ub placed at the central cavity of SIFI, with the substrate HRI located near the DOC2 and KCMF1^N138^ domains, and the fused ubiquitins situated near its corresponding crosslinked regions. Crosslinked residues as determined by mass spectrometry are shown in red and salmon. **e.** Conformational change of the KCMF1^N138^-DOC1-UBR subcomplex upon SIFI bin-ding to HRI^NT^. **f.** SIFI containing a UBR4-subunit lacking its DOC2 domain is impaired in catalyzing ubiquitylation of a mitochondrial presequence. SIFI was purified from WT or *UBR4^ΔDOC2^*cells and incubated with a fluorescently labeled MTS peptide, E1, E2 (UBE2D3 and UBE2A), and ubiquitin. Similar results in n=3 independent experiments. **g.** Deletion of the DOC2 domain in endogenous *UBR4* stabilizes HRI to a similar extent as complete inactivation of *UBR4*. HRI stability was assessed by flow cytometry following GFP-HRI::mCherry reporter expression. Similar results in n=3 independent experiments. **h.** Deletion of the DOC2 domain in UBR4 leads to ISR hyper-activation after mitochondrial stress induced by sodium arsenite. ISR activation was followed by induction of the ATF4 transcription factor by Western blotting. Similar results in n=2 independent experiments. **i.** *UBR4^ΔDOC2^* cells are hyper-sensitive to sodium arsenite-induced stress, dependent on a functional ISR. As indicated, the ISR was inhibited using the small molecule ISRIB. Cell fitness was followed by monitoring cell competition using GFP-labeled WT and mCherry-labeled *UBR4^ΔDOC2^*cells. Each datapoint represents a biological replicate.

SIFI only recognized ABHD10 variants that could be imported into mitochondria (**Extended DataFig. 4d**). Instead of detecting the presequence, SIFI bound ABHD10 that was released from mitochondria during cell lysis and possessed a new N-terminus that resulted from cleavage by the mitochondrial presequence peptidase (**Extended Data** Fig. 4e). SIFI recognized cleaved ABHD10 via the ZZ domain of KCMF1 (**Fig. 3a, b**; **Extended Data** Fig. 3b), which shows structural similarity to a domain in p62 that binds N-degrons (**Extended Data** Fig. 4f) ^27^. AlphaFold models showed that the processed N-termini of ABHD10 fit well into the N-degron binding pocket of KCMF1 (**Fig. 3b**), which was validated by mutation of Asp residues within KCMF1’s N-degron pocket that ablated ABHD10 recognition (**Fig. 3c**). The fortuitous co-purification of ABHD10 thus showed how SIFI detects a set of N-degron targets. We note that DELE1 contains an N-degron that is revealed by stress-induced cleaved by OMA1 ^10^.

**Figure 4:**
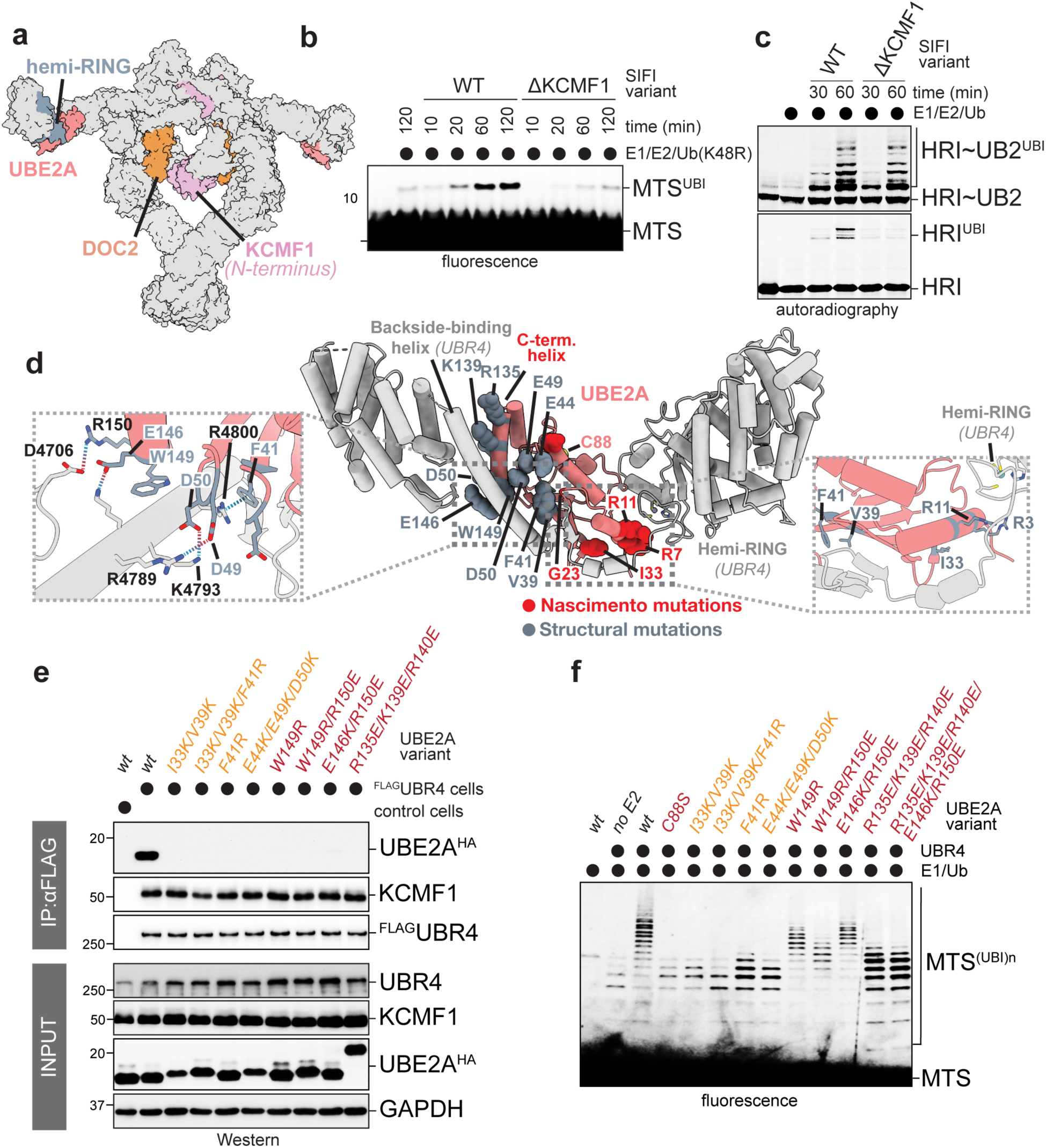
SIFI uses distinct modules for ubiquitin chain initiation and elongation. **a.** Surface representation of SIFI with DOC2 (orange), KCMF1^N138^ (plum), UBE2A (coral), and hemi-RING domain (blue gray) highlighted. **b.** Time-course ubiquitylation assay of a fluorescently labeled pre-sequence peptide by SIFI or SIFI^ΔKCMF1^ indicates that KCMF1 is required for the transfer the first ubiquitin. Reactions were performed in the presence of ubiquitin^K48R^ to prevent chain elongation. Similar results in n=2 independent experiments. **c.** A fusion of two ubiquitin molecules to HRI^NT^ overcomes the defect of SIFI^ΔKCMF1^ in catalyzing ubiquitin chain formation. ^35^S-labeled HRI^NT^∼SUMO (SUMO: for solubility) was incubated with SIFI or SIFI^ΔKCMF1^ and ubiquitylation was followed by autoradiography. **d.** *Middle*: AF2 model of UBE2A bound by the C-terminal region of UBR4 via three interfaces. Mutations introduced to the interfaces are highlighted. *Left*: the interaction network between UBE2A and the backside binding helix of UBR4. *Right*: interactions near the N-terminal helix of UBE2A. **e.** Mutations in UBE2A that disrupt binding to the backside helix or the lock loop of UBR4, respectively, prevent stable integration of UBE2A into SIFI, as revealed by UBR4 affinity-purification and Western blotting using specific antibodies. Experiment performed once and validated by *in-vitro* ubiquitylation. **f.** Mutations shown in **e** impair ubiquitylation of a presequence peptide. Chain initiation occurred via an E2 that co-purified with SIFI.

To determine how SIFI recognizes HRI and unimported mitochondrial proteins, which contain converging degrons ^10^, we incubated it with the degron-containing domain of HRI, HRI^NT^ ^10^. As most E3s bind substrates transiently ^28^, we fused HRI^NT^ to ubiquitin to provide reactive Lys residues for crosslinking to SIFI. Crosslinking combined with mass spectrometry indicated that SIFI recognizes HRI^NT^-Ub at the center of its scaffold, close to the DOC2 domain and C-terminal dimer interface of UBR4 (**Fig. 3d**). The binding of HRI^NT^-Ub caused a substantial shift in the position of the KCMF1 module, while the density ascribed to ABHD10 was strongly reduced (**Fig. 3e**). These results suggested that HRI binds close to UBR4’s DOC2 domain, and hence at a different site than ABHD10, yet both targets occupy overlapping space at the center of the SIFI scaffold.

To assess the role of the DOC2 domain in stress response silencing, we excised it from all *UBR4* loci. We found that deletion of the DOC2 domain in UBR4 impaired the ubiquitylation of mitochondrial presequences (**Fig. 3f**), while it did not affect modification of ABHD10 that is recognized via KCMF1 (**Extended Data** Fig. 5a). Loss of the DOC2 domain stabilized HRI, DELE1, and mitochondrial precursors to a similar extent as complete *UBR4* inactivation (**Fig. 3g**; **Extended Data Fig. 5b**), resulting in increased stress signaling (**Fig. 3h**; **Extended Data** Fig. 5c). SIFI^ΔDOC2^ cells also showed a strong fitness defect when experiencing mitochondrial stress, which was rescued by HRI deletion or treatment with the stress response inhibitor ISRIB (**Fig. 3i**; **Extended Data Fig. 5d**). The DOC2 domains at the center of SIFI therefore play an important role in HRI degradation and stress response silencing.

In addition to ZZ- and DOC2 domains, SIFI contains a UBR box that interacts with N-degrons ^29–32^, yet is not required for stress response silencing ^10^. While the UBR box was not occupied in our structure (**Extended Data** Fig. 6a), its peptide-binding pocket is accessible (**Extended Data Fig. 6b**) and would place targets close to those bound via the ZZ- and DOC2-domains (**Extended Data Fig. 6c**). Notably, the substrate-binding cavity of SIFI shows considerable flexibility in cryo-EM (**Supplementary Movie 3**), which indicates that the E3 ligase could adjust its conformation to its various targets. We conclude that SIFI recognizes its substrates at the center of a flexible scaffold, where they occupy overlapping space and hence compete with HRI to ensure timely stress response silencing.

### SIFI possesses distinct modules for ubiquitin chain initiation and elongation

We noted that substrates bind SIFI at or close to the catalytic ZZ-domain of KCMF1 (**Fig. 4a**). This suggested that KCMF1 helps transfer the first ubiquitin, a reaction that must accommodate the diverse sequence context of Lys residues in SIFI’s many substrates. Loss of KCMF1 indeed impaired attachment of the first ubiquitin (**Fig. 4b**), which was mitigated when we bypassed chain initiation by fusing two ubiquitin molecules to a target (**Fig. 4c**). Once initiation occurred, SIFI^ΔKCMF1^ could build short chains that were connected through the correct K48-linkage (**Extended Data** Fig. 7a). KCMF1 therefore promotes substrate modification, but does not determine the linkage-specificity of SIFI-dependent chain formation.

With KCMF1 supporting initiation, we hypothesized that chain elongation relies on the catalytic modules provided by UBR4. UBR4 only acts with UBE2A or UBE2B ^20^, and co-depletion of these E2s stabilized HRI and increased stress signaling (**Extended Data** Fig. 7b, c). To test if UBR4 drives chain elongation, we assessed the role of E2 enzymes in SIFI-dependent ubiquitylation, making use of the observation that SIFI co-purified with an E2 that could initiate some chain formation if supplied with E1 and ubiquitin. While addition of UBE2D3 increased the abundance of short chains, indicative of a role in initiation, supplementing SIFI with the UBR4-specific UBE2A resulted in selective chain extension (**Extended Data** Fig. 7d).

UBR4 recruits UBE2A through a three-sided embrace that to our knowledge has not been seen for other E3 ligases (**Fig. 4d**). While the hemi-RING of UBR4 binds the N-terminal α-helix of UBE2A, as noted before ^20^, a long UBR4 helix and a short β-sheet together engage the backside of the E2. The helix connects to a UBR4 loop that contacts the C-terminal α-helix of UBE2A (**Fig. 4d**). Mutation of UBE2A residues at each interface prevented integration into SIFI and impaired chain elongation (**Fig. 4e, f**), showing that the E2 embrace is required for SIFI activity.

*UBE2A* mutations that cause the neurodevelopmental Nascimento Syndrome map to each part of the E2 embrace ^33–35^ (**Fig. 4d**): R7 of UBE2A faces the UBR4 hemi-RING; R11 binds the loop that connects the hemi-RING to the backside helix; G23 is next to the backside β-sheet of UBR4; and the C-terminal helix of UBE2A, whose deletion was first noted in Nascimento patients ^35^, engages the UBR4 loop. Mutants in R7, R11, or G23 were impaired in binding to SIFI and catalyzing chain extension (**Extended Data** Fig. 8a, b), which highlights the physiological importance of the E2 embrace and implies that defective stress response silencing contributes to symptoms of Nascimento patients.

### Ubiquitin handover enables linkage-specific ubiquitin chain formation

Since UBE2A lacks linkage specificity ^20,36,37^, additional factors must encode SIFI’s preference for K48 of ubiquitin. SIFI contains a UBL domain between its initiation and elongation modules (**Fig. 5a**). ^1^H-^15^N HSQC NMR spectroscopy showed that this domain binds ubiquitin with a KD of 124μM, an affinity that is sufficient to capture substrate-attached ubiquitin while still enabling processive chain formation (**Fig. 5b**). This interaction relies on UBR4-M4444 and Arg residues mutated in cancer cells, which engage a hydrophobic surface of ubiquitin centered on L8 (**Fig. 5b, c**; **Extended Data Fig. 9a, b**). Strikingly, AlphaFold models showed that the UBL domain orients ubiquitin so that a small movement of the peripheral SIFI arms, as implicated by the flexibility of the UBE2A module in cryo-EM (**Supplementary Movie 4**), is sufficient to present K48 to the active site of UBE2A (**Fig. 5d**). The ε-amino-group of K48 in UBL-bound ubiquitin is guided towards the catalytic Cys88 of UBE2A by Y82 and S120 of the E2, consistent with both residues being essential for chain elongation without affecting UBE2A binding to SIFI (**Extended Data** Fig. 9c, d). These results explain observations that S120 of UBE2A is required for substrate ubiquitylation ^38^, and strongly suggest that the UBL domain captures substrate-attached ubiquitin to hand it over to UBE2A for K48-specific chain formation.

**Figure 5.**
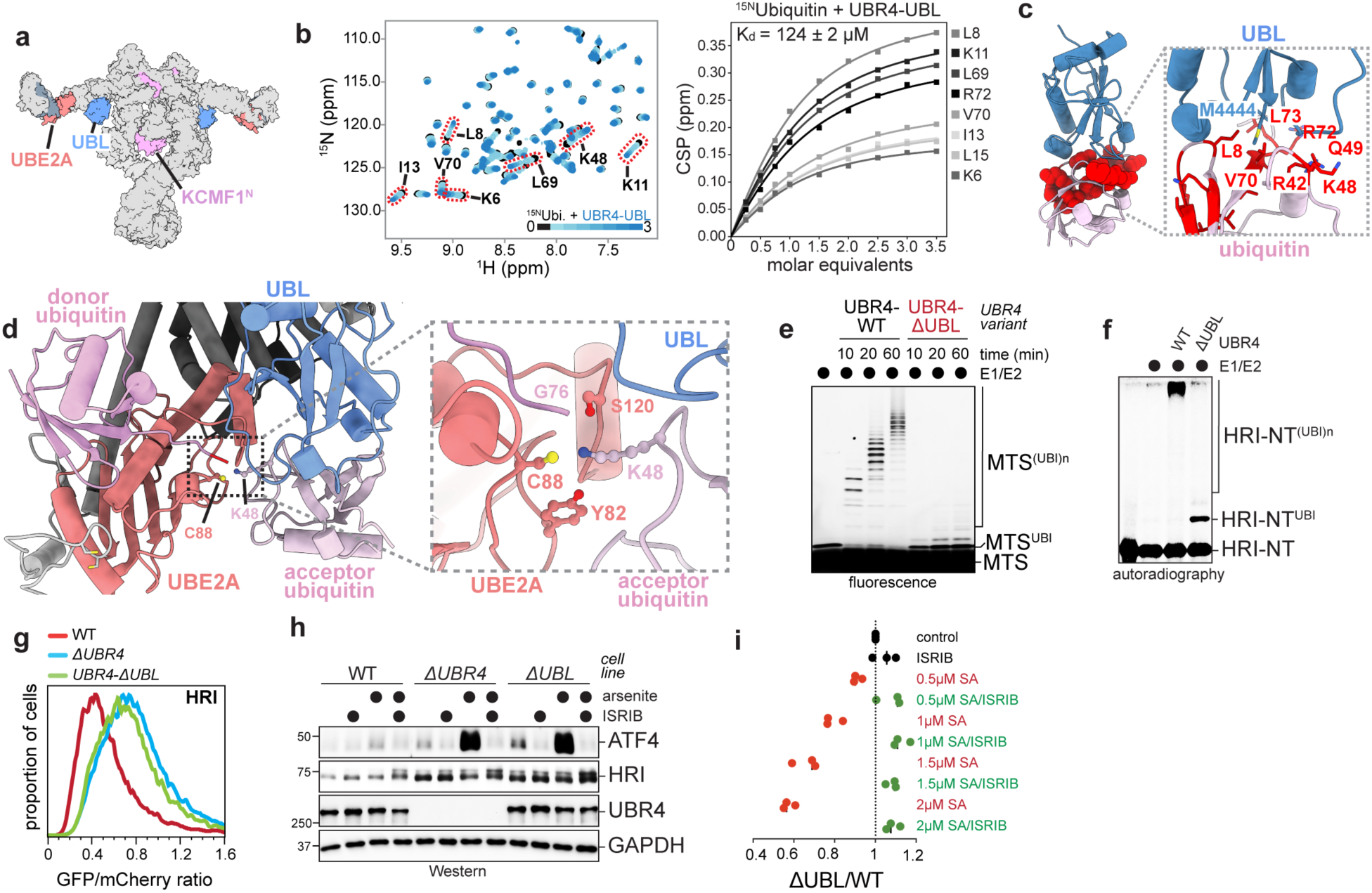
Ubiquitin handover from chain initiation to elongation modules. **a.** Surface representation of SIFI highlighting a ubiquitin-like (UBL) domain located between the KCMF1 initiation site and the UBR4/UBE2A chain elongation center. **b.** *Left*: overlaid ^1^H-^15^N HSQC NMR spectra of ^15^N-labeled ubiquitin titrating unlabeled UBR4 UBL domain. Some highly affected residues are indicated by a red oval. Right: ^1^H-^15^N NMR spectroscopy titration analysis of unlabeled UBL domain and ^15^N-labeled ubiquitin. Binding curves are shown for a subset of residues with well-resolved NMR signals. **c.** Structural model of the UBL-ubiquitin interaction, highlighting interface residues that were highly affected in NMR binding experiments. **d.** Structural model of the catalytic center for ubiquitin transfer. Left: AF3 model illustrating the C-terminal conformational change of UBR4 that brings UBE2A (coral) and the donor Ub (top panel) into close proximity with UBL (blue) and the acceptor Ub (bottom panel). Right: a close-up view of the proposed catalytic center, highlighting G76 of the donor Ub (light pink), K48 of the acceptor Ub (dark pink), and the stabilizing residues S120 and Y82 of UBE2A. **e.** *In vitro* ubiquitylation of a fluorescently labeled presequence peptide shows that SIFI^ΔUBL^ is deficient in ubiquitin chain elongation, but not in initiation. Similar results in n=2 independent experiments. **f.** *In vitro* ubiquitylation of ^35^S-labeled HRI^NT^∼SUMO shows that the UBL domain is required for ubiquitin chain elongation. **g.** Deletion of the UBL domain in *UBR4* stabilizes HRI to the same extent as complete inactivation of *UBR4*. HRI stability was analyzed by flow cytometry using a HRI-GFP::mCherry reporter, as described. Similar results in n=3 independent experiments. **h.** Deletion of the UBL domain in *UBR4* increases activation of the integrated stress response. WT or *UBR4^ΔUBR4^* cells were exposed to sodium arsenite, and stress response activation was monitored via ATF4 accumulation by Western blotting. Similar results in n=2 independent experiments. **i.** UBL deletion in *UBR4* sensitizes cells to sodium arsenite dependent on activation of the integrated stress response. Cell competition assay with sodium arsenite and ISRIB performed as described above. Each datapoint represents a biological replicate.

To assess the physiological importance of the UBL domain, we excised it from *UBR4* and confirmed that SIFI remained intact (**Extended Data** Fig. 10a, b). Strikingly, SIFI^ΔUBL^ was strongly compromised in chain elongation, while transfer of the first ubiquitin was unaffected (**Fig. 5e, f**). HRI, DELE1, and mitochondrial precursors were stabilized by the UBL deletion to the same degree as in *ΔUBR4* cells (**Fig. 5g**; **Extended Data** Fig. 10c), leading to increased stress signaling (**Fig. 5h**; **Extended Data** Fig. 10d). SIFI^ΔUBL^ cells accordingly revealed a strong fitness defect upon stress that was rescued by HRI loss or ISRIB (**Fig. 5i**; **Extended Data** Fig. 10e). Together, these results suggested that the UBL domain hands substrate-attached ubiquitin over to UBE2A for productive chain elongation. Coordination between flexible initiation and linkage-specific elongation modules allows SIFI to efficiently decorate many proteins with a proteolytic ubiquitin tag, a prerequisite for timely and efficient stress response silencing.

## Discussion

Silencing of the integrated stress response requires that a single enzyme, SIFI, can sense stress across cellular scales. Critical to this ability, SIFI processes a large array of mislocalized or cleaved proteins that only accumulate in the cytoplasm during stress and competitively inhibit ubiquitylation of the stress response kinase HRI ^10,13,16,19,29,39^. Our work shows that SIFI can engage such diverse proteins by combining a flexible substrate-modification module with ubiquitin handover to a sterically restricted elongation center (**Fig. 6**).

**Figure 6.**
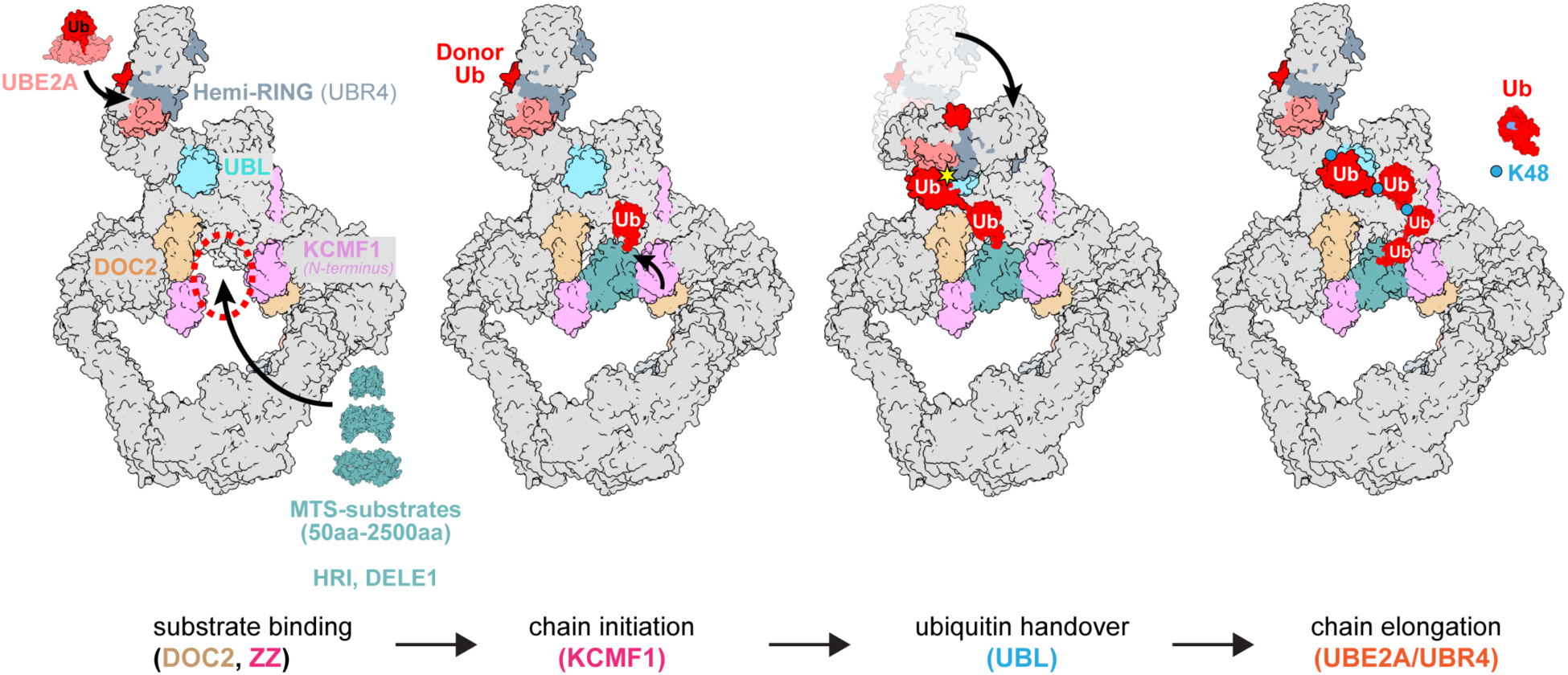
Overview of substrate ubiquitylation by SIFI complex. Substrates bind within the central cavity of SIFI. Ubiquitin chain initiation is mediated by the ZZ-type domain of KCMF1, before a UBL domain hands substrate-attached ubiquitin over to UBE2A for K48-linkage specific chain elongation.

SIFI binds its targets within an easily accessible twisted-ring scaffold. This structure contains at its center substrate-binding and catalytic modules, bringing targets close to active sites responsible for the transfer of the first ubiquitin. These central modules show degrees of flexibility that suggest that they can adapt to targets that range in size from ∼5 to ∼250kDa. However, even substrates with distinct degrons occupy overlapping space on SIFI, which provides a rationale for their competition with HRI as required for timely stress response silencing.

SIFI extends K48-linked chains at separate sites in its peripheral arms. Contrasting the flexibility of the initiation module, SIFI’s elongation sites are built around an E2, UBE2A, that is captured by SIFI on three sides. UBE2A residues at each interface with UBR4 are required for chain extension and mutated in Nascimento Syndrome, showing that the E2 embrace is critical for ubiquitylation. The chain elongation module is therefore sterically restricted. Given that E1 and E3 bind overlapping sites on E2s ^40,41^, it will be interesting to see how UBE2A is recharged with ubiquitin for efficient chain extension.

While ubiquitin chain initiation and elongation can be executed by distinct enzymes ^42,43^, if and how these activities are coordinated is still poorly understood. SIFI directly couples substrate modification to chain elongation by ubiquitin handover through a UBL domain that is tethered to the scaffold by long linkers. The UBL domain orients substrate-attached ubiquitin towards UBE2A so that only K48 is available for chain extension, a feature that requires some movement of the SIFI arms around a hinge close to the UBL domain. Distinct from previously described mechanisms of chain elongation ^44,45^, ubiquitin handover is an efficient solution for enzymes that need to modify many substrates with a specific ubiquitin tag. As ubiquitin-binding domains are found in several quality control E3 ligases ^46–50^, ubiquitin handover might therefore be a more general feature of ubiquitylation machines that must process diverse and often misfolded targets.

Its ability to decorate many proteins with a linkage-specific ubiquitin tag suggests that SIFI should be harnessed for the emerging modality of targeted protein degradation. Our structures point to the ZZ-, DOC2-, and UBR-domains of SIFI as potential binding sites for compounds. As SIFI already counteracts protein aggregation and is highly expressed in brain ^16,51^, it should be particularly tested for applications against neurodegenerative diseases. Our work therefore not only reveals the molecular basis of stress response silencing, but also points to strategies to unlock targeted protein degradation to disease areas of high unmet need.

**Supplementary Movie 1:** Overview of SIFI structure and important domains.

**Supplementary Movie 2:** Movements of the SIFI scaffold around a central hinge.

**Supplementary Movie 3:** Flexibility of the substrate binding cavity of SIFI.

**Supplementary Movie 4:** Movements of the SIFI arms.

## Author contibutions

ZY, DLH and MH performed most experiments, including SIFI purification, cryo-EM analysis, substrate ubiquitylation, and mutant analyses. SW performed NMR analyses. AZ and MJM performed and analyzed crosslink mass spectrometry. TB supported biochemical analyses of SIFI. MR helped plan and interpret experiments, and ZY, DLH, MH, SW and MR wrote the manuscript.

## Acknowledgements

We thank all members of our laboratory for enthusiastic support and many suggestions for this work. We thank Drs. Dan Toso, Ravindra Thakkar, Paul Tobias, and Cal-Cryo cryo-EM facility at QB3-Berkeley for assistance on cryo-EM data collection. We thank Drs. Hasan Celik, Raynald Giovine, and Pines Magnetic Resonance Center’s Core NMR Facility (University of Washington) for spectroscopic assistance, and we are grateful to Dr. Peter Brzovic (University of Washington) for sharing ubiquitin NMR chemical shifts. The instruments used in this work were supported by the PMRC Core. We thank Dr. Majid Ghassemian and the UCSD Biomolecular and proteomics mass spectrometry facility for assistance in running the mass spec samples. DLH was funded by an HHMI Helen Hay Whitney Fellowship. MH is funded by the Deutsche Forschungsgemeinschaft (DFG, German Research Foundation; HE 9330/1-1). This work was supported by the G. Harold and Leila Y. Mathers Charitable Foundation and the Howard Hughes Medical Institute. MR is an Investigator of the Howard Hughes Medical Institute.

## Material and Methods

### Data reporting

No statistical methods were used to predetermine sample size. The experiments were not randomized and the investigators were not blinded to allocation during experiments and outcome assessment.

### Mammalian cell culture

HEK293T were maintained in DMEM + Glutamax (Gibco 10566-016) plus 10% fetal bovine serum (VWR 89510-186). HEK293T cells that were adapted to growth in suspension were cultured in FreeStyle 293 Expression medium (Gibco 12338026) with 1% fetal bovine serum. Expi293F cells used for transient transfection for protein purification were grown in Expi293 Expression medium (A1435101, Gibco). All cell lines were purchased directly from the UC Berkeley Cell Culture Facility, authenticated by short tandem repeat analysis and were routinely tested for mycoplasma contamination using the Mycoplasma PCR Detection Kit (abmGood, G238). All cell lines tested negative for mycoplasma.

Plasmid transfections were performed using polyethylenimine (PEI, Polysciences 23966-1) at a 1:6 ratio of DNA (in μg) to PEI (in μl at a 1 mg/ml stock concentration), Lipofectamine 3000 transfection reagent (ThermoFisher, L3000008) or Expifectamine per manufacturer’s instructions or ExpiFectamine 293 (ThermoFisher, A14524). Lentiviruses were produced in HEK293T cells by cotransfection of lentiviral and packaging plasmids using PEI. Virus containing supernatants were collected 48h and 72h after transfection, supernatants were spun down aliquoted, and stored at −80°C for later use. For lentiviral transduction, 10^5^ cells were seeded into 24-well plates and subjected to centrifugation for 45 minutes at 1000xg after addition of lentiviral particles and 6μg/ml polybrene (Sigma-Aldrich, TR-1003). HEK293T transduced cells were drug-selected 24h post infection with the following drug concentrations when applicable: puromycin (1μg/ml, Sigma-Aldrich, P8833), blasticidin (7.5μg/ml, ThermoFisher, A1113903).

### Plasmids

The list of all constructs used in this study is provided in Supplementary Table 1. Most cloning was performed using Gibson assembly with the HIFI DNA Assembly master mix (NEB, E2621L).

### Generation of CRISPR-cas9 genome edited cell lines

All CRISPR-cas9 edited cell lines used in this study were generated from HEK293T cells. sgRNA sequences were designed using the on-line resource provided by IDT. DNA oligonucleotides for sgRNA and their complementary sequence were phosphorylated (NEB, M0201), annealed and ligated (NEB, M0202) into pX330 (Addgene, #42230). HEK293T cells were cultured in a 6-well plate and transfected at 50% confluence with 2μg of px330 plasmids (and 1μl of 10μM single stranded donor oligo when applicable) using Mirus TransIT-293 Transfection reagent (Mirus, MIR2705). 48h post transfection, individual clones were expanded in 96-well plates. Homozygous clones were screened by PCR and DNA sequencing and confirmed by western blotting when applicable.

HEK293T ^FLAG^UBR4, ΔUBR4, ^FLAG^UBR4 ΔKCMF1 cells were generated previously ^10^. For the generation of N-terminally Twin-strep UBR4 cells (^TwinStrep^UBR4), we used the following sgRNA: 5’-gcggaagatggcgacgagcgg -3’ and ssODN 5’-ccggtggcaagcccccggagggagccgcagtagtacgacggaagatgagcgcctggagtcaccctcagtttgagaaaggcggaggtagcggaggtggctctggcggaagcgcctggtcacacccacagttcgagaagggcggaggtagcgcgacgagcggcggcgaagaggcggcggcagcggctccggcgccggg -3’. UBR4^ΔUBL^(Δ4342-4434), UBR4^ΔDOC2^ (Δ3538-3721) were generated in the ^FLAG^UBR4 background, with the following protospacer sequences that created in-frame deletions: UBR4^ΔUBL^:

5’-ccttgttccatggacactcg -3’ and 5’- cattagtcagatgcctccaa -3’; UBR4^ΔDOC2^ : 5’-cgggttattacacaccaggc -3’ and 5’- atgggatccactgcacagca -3’ and the following sODN to repair: 5’- catgctaagactggtttcttccttagcactttgtctggcttagtggagtttgatggctattacctggagagcgatccctgcctcgtgtgtaataacggcagcagcgcagtggatcctattgagaatgaagaagaccggaagaaggtgaggccagatctggcctagactcagggctgtggccttgatctggactttgggca-3’.

### Protein expression and purification

To purify the endogenous SIFI complex, 293T cells with the *UBR4* gene endogenously tagged with a FLAG or Strep tag were harvested from 150mm culture dishes or adapted suspension cells. To purify SIFI complex through affinity-tagged KCMF1, TwinStrep II-tagged KCMF1 was transiently expressed in Expi293F cells for 48 hours before harvesting. Cells were lysed in cell lysis buffer containing 40mM HEPES pH7.5, 150mM NaCl, 1mM DTT, 0.1% NP-40, Benzonase nuclease (Millipore Sigma, 70746-4), proteasome inhibitor carfilzomib, and Roche cOmplete protease inhibitor cocktail (Sigma, 11836145001). Cells were lysed for 20 minutes and homogenized using a Dounce homogenizer. Cell lysate was cleared by a 4,000 xg centrifuge for 10 minutes followed by 36,000 xg centrifuge for 40 minutes. Supernatant solution was subsequently collected and incubated with M2 FLAG resin (Sigma-Aldrich, A2220) or Strep-Tactin XT 4Flow resin (IBA Lifesciences GmbH, 16674714) for 4-6 hours at 4°C. Affinity resin was washed thoroughly with wash buffer (150mM HEPES pH 7.5, 150mM NaCl, 1mM DTT), then eluted with elution buffer (500 μg/mL 3xFLAG peptide (Sigma, F4799) for M2 FLAG resin, 50mM biotin (IBA Lifesciences GmbH, 21016005) for Strep-Tactin XT 4Flow resin). Eluted proteins further purified with a Superose 6 10/300 size-exclusion chromatography (SEC) column (Cytiva, 17517201).

### Cryo-EM sample preparation

Purified SIFI complex, SIFI-substrate mixture, or crosslinked SIFI-substrate mixture samples, were concentrated to 2-4 mg/mL concentration for cryo-preservation. Concentrated samples were mixed with a final concentration of 0.02% (w/v) fluorinated octylmaltoside (Anatrace) immediately before cryo-freezing to prevent protein denaturation at the air-water interface. 2.6 μL of the sample was applied to a glow-discharged 300-mesh Quantifoil R1.2/1.3 grid and incubated for 15 seconds before being blotted and plunge vitrified in liquid ethane, which was cool-protected by liquid nitrogen. Grid freezing was performed using a Mark IV Vitrobot (ThermoFisher Scientific) system operating at 12°C and 100% humidity.

### Cryo-EM data collection and processing

Cryo-EM data collections were carried out using a 300kV Titan Krios G3i or G2 electron microscope (ThermoFisher Scientific) equipped with a BIO Quantum energy filter (slit width 20 eV). Data were collected using SerialEM software ^52^ at a nominal 105,000x magnification with a pixel size of 1.05 Å/pixel (G3i) or 0.83 Å/pixel (G2). Movies were recorded using a 6k x 4k Gatan K3 Direct Electron Detector operating at super-resolution CDS mode. Each movie was composed of 40 subframes with a total dose of 60 e^-^/Å^2^, resulting in 1.5 e^-^/Å^2^ dose rate. A total of >6,000 movies were recorded for each sample. Data processing including motion correction, CTF estimation, particle picking, 2D class averaging, and 3D refinement were done using cryoSPARC v4.3 workflow ^53^. All movies were 2x binned and patch motion-corrected. After particle picking and several iterations of 2D class averaging, initial 3D volume was calculated using *ab initio* 3D reconstruction. Selected particles were then used for non-uniform 3D refinements ^54^ Protein motion and flexibility was calculated using cryoSPARC 3D Flexible Refinement ^55^ (3DFlex).

### Model building and structural analysis

Coordinates for the N- and C-terminal halves of SIFI complex were built separately into their respective cryo-EM maps. Coordinates for regions of the EM maps with higher local resolutions, e.g. the UBR4 α-helical Armadillo repeats, were manually built into the map, whereas for regions with lower local resolutions (∼5-8 Å), AlphaFold2 ^56^ models were used and protein secondary structures, including α-helices and β-strands, were fitted into the EM density. For regions with very low resolutions (< 8 Å), e.g. UBR4 WD40 domain, UBR, DOC1, the N-terminal domain of KCMF1, UBL domain, and hemi-RING/UBE2A, AlphaFold2 models were fitted into the low-resolution density in ChimeraX without refining amino acid side chains found in these domains. Certain low-resolution models, such as the UBL domain of UBR4, the UBE2A-binding domains, and the N-terminal domain of KCMF1, were validated through other biochemical assays such as NMR, binding assays, and *in vitro* ubiquitylation assays. Coordinates of the medium to high resolution regions were refined with multiple iterations of PHENIX real-space refinements ^57^ and manual refinement in Coot ^58^. Buried surface area (BSA) of protein-protein interface was calculated using PDBePISA ^59^ ith a 1.4Å probe. Potential hydrogen bonds were assigned using the geometry criteria of <3.5 Å separation distance and >90° A-D-H angle. A maximum distance of 4.0 Å was allowed for a potential van der Waals interaction. Structures were further analyzed, and structural presentations were prepared using PyMOL v2 (Schödinger LLC) and ChimeraX^60,61^.

### Mass spectrometry

Mass spectrometry was performed on immunoprecipitates prepared from forty 15-cm plates of endogenously ^FLAG^UBR4 and ^FLAG^UBR4 ΔKCMF1 HEK 293T cell lines or twenty 15-cm plates of WT or ΔUBR4 HEK293T cells transfected with KCMF1-3xFLAG DNA. Cells were lysed in lysis buffer (40 mM HEPES, pH 7.5, 150 mM NaCl, 0.2% Nonidet P-40, benzonase (Sigma-Aldrich, E1014) and 1× cOmplete protease inhibitor cocktail (Roche, 11836170001), 1xPMSF), lysed extracts were clarified by centrifugation at 21,000xg, and bound to anti-FLAG M2 affinity resin (Sigma-Aldrich, A2220) for 2 hours rotating at 4°C. IPs were then washed 4x and eluted 3x at 30°C with 0.5 mg/ml of 3×FLAG peptide (Sigma, F4799) buffered in 1×PBS plus 0.1% Triton X-100. Elutions were pooled and precipitated overnight at 4°C with 20% trichloroacetic acid. Spun down pellets were washed 3x with an ice-cold acetone/0.1 N HCl solution, dried, resolubilized in 8 M urea buffered in 100 mM Tris 8.5, reduced with TCEP, at a final concentration of 5 mM, (Sigma-Aldrich, C4706) for 20 min, alkylated with iodoacetamide, at a final concentration of 10 mM, (ThermoFisher, A39271) for 15 min, diluted four-fold with 100 mM Tris-HCl 8.5, and digested with 0.5 mg/ml of trypsin (Promega,v5111) supplemented with CaCl2 (at a final concentration of 1 mM) overnight at 37°C. Trypsin-digested samples were submitted to the UC Berkeley Vincent J. Coates Proteomics/Mass Spectrometry Laboratory for analysis. Peptides were processed using multidimensional protein identification technology (MudPIT) and ran on a LTQ XL linear ion trap mass spectrometer. To identify high confidence interactors, CompPASS analysis was performed against mass spectrometry results from unrelated FLAG immunoprecipitates performed in our laboratory. KCMF1, ABHD10 and NIPSNAP3A were the only three proteins with more than 10 spectral counts and showing a ten-fold or more reduction in ^FLAG^UBR4 ΔKCMF1 compared to ^FLAG^UBR4.

^FLAG^UBR4 and ^FLAG^UBR4^ΔUBL^ mass spectrometry experiments (**Extended Data** Fig. 10a) were performed from 20 15-cm plates lysed in lysis buffer (40 mM HEPES, pH 7.5, 150 mM NaCl, 0.2% Nonidet P-40, benzonase (Sigma-Aldrich, E1014) and 1× cOmplete protease inhibitor cocktail (Roche, 11836170001), 1xPMSF), clarified by centrifugation at 21,000xg, and bound to anti-FLAG M2 affinity resin (Sigma-Aldrich, A2220) for 2 hours rotating at 4°C. IPs were then washed 4x with (40 mM HEPES, pH 7.5, 150 mM NaCl, 0.2% Nonidet P-40) followed by 3 washes in PBS, all remaining liquid was removed with a crushed gel tip and the beads were flash frozen in liquid nitrogen. Further sample processing was performed at UC San Diego Proteomics Facility where protein samples were diluted in TNE (50 mM Tris pH 8.0, 100 mM NaCl, 1 mM EDTA) buffer. RapiGest SF reagent (Waters Corp.) was added to the mix to a final concentration of 0.1% and samples were boiled for 5 min. TCEP was added to 1 mM (final concentration) and the samples were incubated at 37°C for 30 min. Subsequently, the samples were carboxymethylated with 0.5 mg/ml of iodoacetamide for 30 min at 37°C followed by neutralization with 2 mM TCEP (final concentration). Proteins samples were then digested with trypsin (trypsin:protein ratio - 1:50) overnight at 37°C. RapiGest was degraded and removed by treating the samples with 250 mM HCl at 37°C for 1 h followed by centrifugation at 14000 rpm for 30 min at 4°C. The soluble fraction was then added to a new tube and the peptides were extracted and desalted using C18 desalting columns (Thermo Scientific, PI-87782). Peptides were quantified using BCA assay and a total of 1 ug of peptides were injected for LC-MS analysis. Trypsin-digested peptides were analyzed by ultra high pressure liquid chromatography (UPLC) coupled with tandem mass spectroscopy (LC-MS/MS) using nano-spray ionization. The nanospray ionization experiments were performed using a TimsTOF 2 pro hybrid mass spectrometer (Bruker) interfaced with nano-scale reversed-phase UPLC (EVOSEP ONE). Evosep method of 30 SPD (samples per day) was utilized using a 10 cm × 150 μm reverse-phase column packed with 1.5 μm C18-beads (PepSep, Bruker) at 58 °C. The analytical columns were connected with a fused silica ID emitter (10 μm ID; Bruker Daltonics) inside a nanoelectrospray ion source (Captive spray source; Bruker). The mobile phases comprised 0.1% FA as solution A and 0.1% FA/99.9% ACN as solution B. The mass spectrometry settings for the TimsTOF Pro 2 are as follows: The dia-PASEF method for proteomics. The values for mobility-dependent collision energy was set to 10 eV. No merging of TIMS scans was performed. The ion mobility (IM) was set between 0.85 (1/k0) and 1.3 (1/k0) with a ramp time of 100 ms. Each method includes one IM windows per dia-PASEF scan with variable isolation window at 20 amu segments. 34 PASEF MS/MS scans were triggered per cycle (1.38 s) with a maximum of seven precursors per mobilogram. Precursor ions in an *m*/*z* range between 100 and 1700 with charge states ≥3+ and ≤8+ were selected for fragmentation. Protein identification and label free quantification was carried out using Spectronaut 18.0 (Biognosys). The values obtained with DIA analysis for the ^FLAG^UBR4 and ^FLAG^UBR4^ΔUBL^ values are represented for a subset of interactors that were found significantly enriched over the 293T background control samples and were previously validated.

### Protein crosslinking

Cross-linking reactions were performed in 40 mM HEPES pH 7.4 150 mM NaCl 1 mM DTT buffer containing 3 mg/mL total protein. BS3 (bis(sulfosuccinimidyl)suberate) (ThermoFisher, A39266) dissolved in water was added to a final concentration of 0.5 mM to initiate cross-linking. Cross-linking was performed at room temperature (RT) for 30 minutes prior to quenching by addition of 100 mM final concentration of ammonium bicarbonate. Cross-linked samples were reduced with 10 mM final concentration of TCEP for 1 hour at 37°C, alkylated with 15 mM final concentration of iodoacetamide at RT in the dark for 30 mins followed by addition of 15 mM final concentration of DTT. Sequencing Grade Modified Trypsin (Promega, V5111) was added at an enzyme:substrate ratio of 1:15. Tryptic digestion was performed for 6 hours at 37°C prior to acidification by addition of HCl to a final concentration of 250 mM. Digested samples were spun in a microfuge at 21,000 g for 10 minutes and the resulting supernatant transferred to an autosampler vial and stored at -80°C prior to analysis by liquid chromatography with tandem mass spectrometry (LC-MS/MS).

### Mass Spectrometry of crosslinked samples

Mass spectrometry and data analysis were based on previously described methods ^62^. Sample digest (5 µL) was loaded onto a PepMap Neo Trap Cartridge (Thermo Scientific174500). The trap was brought online with a Bruker PepSep C18 15 cm x 150 µm, 1.9 µm column (Thermo Scientific,1893471) connected to a 5 cm x 20 µm ID Sharp Singularity Fossil Ion Tech tapered tip mounted in a custom constructed microspray source. Peptides were eluted from the column at 0.8 µL/min using a 90 minute acetonitrile gradient. An Orbitrap Exploris 480 (Thermo Fisher Scientific) was used to perform MS in data dependent acquisition (DDA) mode using the following parameters. Three methods were used, differing only by MS/MS resolution. The resolution at *m/z* 200 for MS was 60,000. MS/MS resolution was 15,000 or 30,000 or 60,000. Cycle time was 2 seconds between MS scans. MS scan range was *m/z* 400–1,600. The automatic gain control was set to standard for MS and MS/MS, and the maximum injection times were set to auto. MS/MS spectra were acquired using an isolation width of 2 *m/z* and a normalized collision energy of 27. MS/MS acquisitions included +3 to +6 precursor ions and undefined precursor charge states were excluded. Dynamic exclusion was set at 10 s. All spectra were collected in centroid mode.

Acquired spectra were converted to mzML format using ProteoWizard’s msConvert ^63^. Crosslinks were identified from DDA data using the Kojak ^64^ search algorithm with post-processing using Percolator ^65^ and cross-link data visualization using the ProXL web application ^66^. All data were filtered at a false discovery rate (FDR) of 5% at the peptide level unless otherwise stated.

### Growth competition assays

HEK293T, ΔUBR4, ΔDOC2 and ΔUBL cells were transduced to express either GFP or mCherry, respectively using the lentiviral pLVX-GFP-P2A-Blasticidin or pLVX-mCherry-P2A-Blasticidin vector as described previously ^10^.

For drug competition assays, 5x10^4^ wt-GFP and 5x10^4^ ΔUBR4-mCherry or UBR4 domain mutants were mixed in 6-well plates. The next day, indicated concentrations of sodium arsenite (Ricca Chemical, 714216), were added for 72 hours. The ratio of mCherry+/GFP+ cells was determined on a BD LSRFortessa instrument, analyzed with FlowJo 10.8.1 and normalized to the untreated sample. Gating strategies for flow cytometry analysis are shown in Supplementary Fig. 1.

### Drug treatments

For 3 days growth competition experiments with drug treated cells, we used the following drug concentrations: 0.5-2 μM sodium arsenite (Ricca Chemical, 714216). For overnight drug treatments we used 5 μM sodium arsenite, or otherwise indicated in the figure legends. To inhibit the proteasome, we used 2 μM Carfilzomib (Selleck Chemicals, S2853) for 6 h. ISRIB (Sigma-Aldrich, SML0843) was used at a concentration of 200nM.

### Protein stability reporter assay

The pCS2+-degron-GFP-IRES-mCherry reporter constructs (ISR, HRI, and DELE1) were generated as described ^10^ and are listed Supplementary Table 1. Protein stability reporter assays were performed as described previously ^10^. Cells were analyzed on either a BD Bioscience LSR Fortessa or a LSR Fortessa X20 and the GFP/mCherry ratio was analyzed using FlowJo. Gating strategies for flow cytometry analysis are shown in Supplementary Fig. 1.

### Western blotting

For western blot analysis of whole cell lysates, cells were harvested at indicated time points by washing in PBS, pelleting and snap freezing. Cells were lysed in lysis buffer (150mM NaCl, 50mM HEPES pH 7.5, 1% NP-40 substitute) supplemented with Roche cOmplete Protease Inhibitor Cocktail (Sigma, 11836145001), PhosSTOP™ Phosphatase Inhibitor Cocktail (Roche, 4906837001), carfilzomib (2μM) and benzonase (EMD Millipore, 70746-4) on ice. Samples were then normalized to protein concentration with Pierce 660nm Protein Assay Reagent (Thermo Fisher, 22660). 2x urea sample buffer (120 mM Tris pH 6.8, 4% SDS, 4 M urea, 20% glycerol, bromophenol blue) was then added to the samples. SDS-PAGE and immunoblotting was performed with the indicated antibodies. Images were captured on a ProteinSimple FluorChem M device.

### Small-scale Immunoprecipitations

Cells were harvested after washing in PBS, pelleted and snap frozen. Frozen pellets were resuspended in lysis buffer (40mM HEPES 7.5, 150mM NaCl, 0.1% NP-40 (Nonidet P-40), with Roche cOmplete Protease Inhibitor Cocktail (Sigma-Aldrich, 11873580001), carfilzomib (2μM, Selleckchem, S2853) and benzonase (EMD Millipore, 70746-4). Lysates were incubated for 30 minutes on ice and cleared by centrifugation for 20min at 21000xg, 4°C. Supernatants were normalized to volume and protein concentration. 5% of the sample was removed as an input and the sample was added to equilibrated ANTI-FLAG-M2 Affinity Agarose Gel slurry (Sigma-Aldrich, A2220) and rotated for 1-2 hours at 4°C. Beads were washed three times in wash buffer (40mM HEPES 7.5, 150mM NaCl, 0.1% NP40) and eluted with 2x urea sample buffer. SDS-PAGE and immunoblotting was performed with the indicated antibodies. Images were captured on a ProteinSimple FluorChem M device.

For immunoprecipitations performed from cells lysed in digitonin lysis buffer, cells were harvested after washing in PBS and immediately lysed (never frozen) in digitonin lysis buffer (40mM HEPES 7.5, 150mM NaCl, 50μg/ml digitonin (Sigma-Aldrich, D141), with Roche cOmplete Protease Inhibitor Cocktail (Sigma-Aldrich, 11873580001), carfilzomib (2μM, Selleckchem, S2853) for 10 min rotating at 4C. Lysed cells were then spun down at 2000xg for 5 min at 4C and the supernatant was collected, 5% input was removed. The supernatant was added to equilibrated ANTI-FLAG-M2 Affinity Agarose Gel slurry (Sigma-Aldrich, A2220) and rotated for 1-2 hours at 4°C. Following wash and elution steps were the same as described above for NP-40 lysis. The pellet obtained after centrifugation of the digitonin lysed cells was also subsequently lysed in NP-40 lysis buffer as described above to break open mitochondria and sample was collected after centrifugation at 21000xg as the NP-40 fraction.

### Antibodies

Following antibodies were used for immunoblot analyses: anti-Flag (mouse, Clone M2, Sigma-Aldrich, F1804, dilution 1:1000), anti-Flag (rabbit, Cell Signaling Technology (CST), 2368, dilution 1:1000), anti-HA-Tag (rabbit, C29F4, CST, 3724, dilution 1:1000), anti-strep (strepMAB-Classic, 2-1507-001, iba lifesciences, dilution 1:10000), anti-GAPDH (rabbit, D16H11, CST, 5174, dilution 1:1000), anti-HSP90β (rabbit, D3F2, CST, 7411, dilution 1:1000), anti-α Tubulin (mouse, DM1A, Calbiochem, CP06, dilution 1:1000), anti-UBR4/p600 (rabbit, A302, Bethyl, A302-279A, dilution 1:1000), anti-UBE2A/B (mouse, G-9, Santa Cruz, sc-365507, dilution 1:150), anti-ATF4 (rabbit, D4B8, CST, 11815S, dilution 1:1000), anti-EIF2AK1 (rabbit, Proteintech, 20499-1-AP, dilution 1:1000), anti-KCMF1 (rabbit, Sigma, HPA030383, dilution 1:1000), anti-NIPSNAP3A (rabbit, ThermoFisher, PA5-20657, dilution 1:1000), anti-Ubiquitin (rabbit, Cell Signaling Technology (CST), 43124, dilution 1:1000), anti-ABHD10 (rabbit, ThermoFisher, PA5-103553, dilution 1:1000), goat anti-rabbit IgG (H+L) HRP (Vector Laboratories, PI-1000, dilution 1:5000), Sheep anti-mouse IgG (H+L) HRP (Sigma, A5906, dilution 1:5000), goat anti-mouse IgG light chain specific HRP conjugated (Jackson Immunoresearch, 115-035-174, dilution 1:5000).

### In vitro transcription/translation of substrates

In vitro synthesized substrates were all cloned into pCS2 vectors containing a SP6 promoter (Supplementary Table 1) and generated using Wheat Germ Extract (Promega, L3260) as previously described ^10^.

### In vitro ubiquitylation assays

For in vitro ubiquitylations, human SIFI complex was purified using an endogenous ^FLAG^UBR4 HEK 293T cell line. Each in vitro ubiquitylation reaction required material from 2.5 15-cm plates of ^FLAG^UBR4, ^FLAG^UBR4 ^ΔUBL,^ or ^FLAG^UBR4 ^ΔDOC2^. For ^FLAG^UBR4 ΔKCMF1 we used 10 15-cm plates per reaction. Frozen cell pellets were lysed at 4°C for 30 min in 1 ml of lysis buffer per 10 15-cm plates (40 mM HEPES, pH 7.5, 5 mM KCl, 150 mM NaCl, 0.1% Nonidet P-40, 1mM DTT, 1X cOmplete protease inhibitor cocktail, 2 μM carfilzomib and 4 μl of benzonase per 10 15-cm plates). Lysed extracts were pelleted at 21,000xg to remove cellular debris and the clarified lysate was bound to anti-FLAG M2 resin (20 μl of slurry per 2.5 15-cm plates of material) for 2h rotating at 4°C. UBR4-coupled beads were washed 2x with (40 mM HEPES, pH 7.5, 5 mM KCl, 150 mM NaCl, 0.1% Nonidet P-40, 1mM DTT) and 2x without detergent (40 mM HEPES, pH 7.5, 5 mM KCl, 150 mM NaCl, 1mM DTT), all liquid was removed from the beads using a crushed gel loading tip prior to addition of the in vitro ubiquitylation reaction.

In vitro ubiquitylation assays were performed in a 10 μl reaction volume: 0.5 μl of 10 μM E1 (250 nM final), 0.5 μl of 50 μM Ube2A (2.5 μM final), 0.5 μl of 50 μM Ube2D3 (2.5 μM final), 1 μl of 10 mg ml-1 ubiquitin (1 mg ml^-1^ final) (R&D Systems, U-100H), 0.5 μl of 200 mM DTT, 1.5 μl of energy mix (150 mM creatine phosphate (Sigma-Aldrich, 10621714001-5G), 20 mM ATP, 20 mM MgCl2, pH to 7.5 with NaHCO3), 1 μl of 10x ubiquitylation assay buffer (250 mM Tris 7.5, 500 mM NaCl, and 100 mM MgCl2), 0.5 μl of 1mg/ml TUBEs (Tandem Ubiquitin Binding Entities) were pre-mixed and added to 10 μl of UBR4-coupled bed resin. 3 μl of in vitro translated substrate, 1 μl of 100μM TAMRA-labeled peptide were added to the reactions. In the case of ABHD10-3xHA immunoprecipitated with ^FLAG^UBR4 or KCMF1-3xFLAG (and mutants) immunoprecipitates no additional substrate was added. PBS was added to reach final volume of 10 μl. Peptide sequences used in this study are summarized in Supplementary Table 2. Reactions were performed at 30°C with shaking for 2 h or as indicated in the respective timecourse. Reactions were stopped by adding 2X urea sample buffer and resolved on SDS-PAGE gels prior to autoradiography in the case of radio-labeled substrates. MTS-TAMRA peptide ubiquitylations were run on 4-20% gradient gels (ThermoFisher, EC6026BOX) and imaged on a ProteinSimple Fluorchem M imager. ABHD10-3xHA or KCMF1-3xFLAG ubiquitylations were visualized by western blotting with anti-HA or anti-ubiquitin antibodies, respectively. Commercially available recombinant human ubiquitin mutants used in this study included (R&D Systems, UM-K48R, UM-K480, UM-NOK). L8A ubiquitin was purified as described previously ^44^. E1 enzyme UBA1 was purified as described in ^67^. Ube2A and Ube2A mutants, Ube2D3 and Tube recombinant proteins were purified as previously described ^10^.

### Protein purification for NMR analysis

For NMR isotopic labeling, wild-type untagged ubiquitin was expressed in E. coli BL21 (DE3) grown in M9 minimal media supplemented with ^15^N-ammonium chloride (Cambridge Isotope Labs). Ubiquitin was purified via cation exchange followed by size exclusion chromatography (SEC) using a Superdex 75 column (GE Healthcare) equilibrated in NMR buffer (25 mM NaPi pH 7.0, 150 mM NaCl, 1 mM TCEP) as previously described ^68^. 6xHis-tagged UBR4-UBL (residues 4340-4463; UniProt Q5T4S7-1) variants were expressed in E. coli BL21 (DE3) and purified via Ni^2+^-NTA affinity chromatography using a Cytiva HisTrap FF crude column according to manufacturer’s instructions followed by SEC using a Superdex 75 column equilibrated in NMR buffer.

### NMR

All NMR samples were assembled with 200mM ^15^N-labeled ubiquitin and indicated amounts of unlabeled UBR4-UBL in NMR buffer supplemented with 10% deuterium oxide. ^1^H-^15^N hetero-nuclear single quantum coherence (HSCQ) spectra were collected on a Bruker Ascend 500mHz magnet equipped with a Bruker Avance IV NEO console and a 5mm BBO Prodigy CryoProbe. Data were transformed and phased using Bruker TopSpin (4.3) and plotted in NMRViewJ. Combined amide (NH) chemical shift perturbations (CSP) were calculated using the following equation: ΔδNH (ppm) = sqrt [ΔδH^2^ + (ΔδN/5)^2^]. NMR CSP titration data were fitted to the standard equation for a 1:1 binding equilibrium to derive a dissociation constant (Kd) for the ubiquitin-UBL interaction: ΔδNH (ppm) = Δδmax (([P]t + [L]t + Kd) – [([P]t + [L]t + Kd)^2^ – 4[P]t[L]t]^1/2^)/ 2[P]t where [L]t and [P]t are the concentrations of ligand (UBR4-UBL) and protein (ubiquitin), respectively ^69^. A subset of NMR signals with substantial and well-resolved chemical shifts were used for Kd calculation. Data were plotted and fit to the binding equilibrium equation using a custom python script. Plotted curves were fit to individual residues, while the reported Kd represents a global fit to all plotted residues with the standard deviation derived from the residual sum of squares.

### Software and code for data analysis

The following freely/commercially available softwares/codes were used to analyze data: FlowJo (v.10.8.1), GraphPad Prism (v.9), NMRViewJ (v9.2.0-b27), Bruker TopSpin (v.4.3.0), cryoSPARC (v.4.3), SerialEM. (v.4.1), 3DFlex, AlphaFold (v.2; v.3), PHENIX (v.1.21.1-5286), Coot (v.0.9.8.92), PDBePISA (v1.52), Chimera (v.1.17.1), ChimeraX (v.1.8), PyMOL (v.2.5.5), Spectronaut (18.0),

---

**Extended Data Figure 1:**
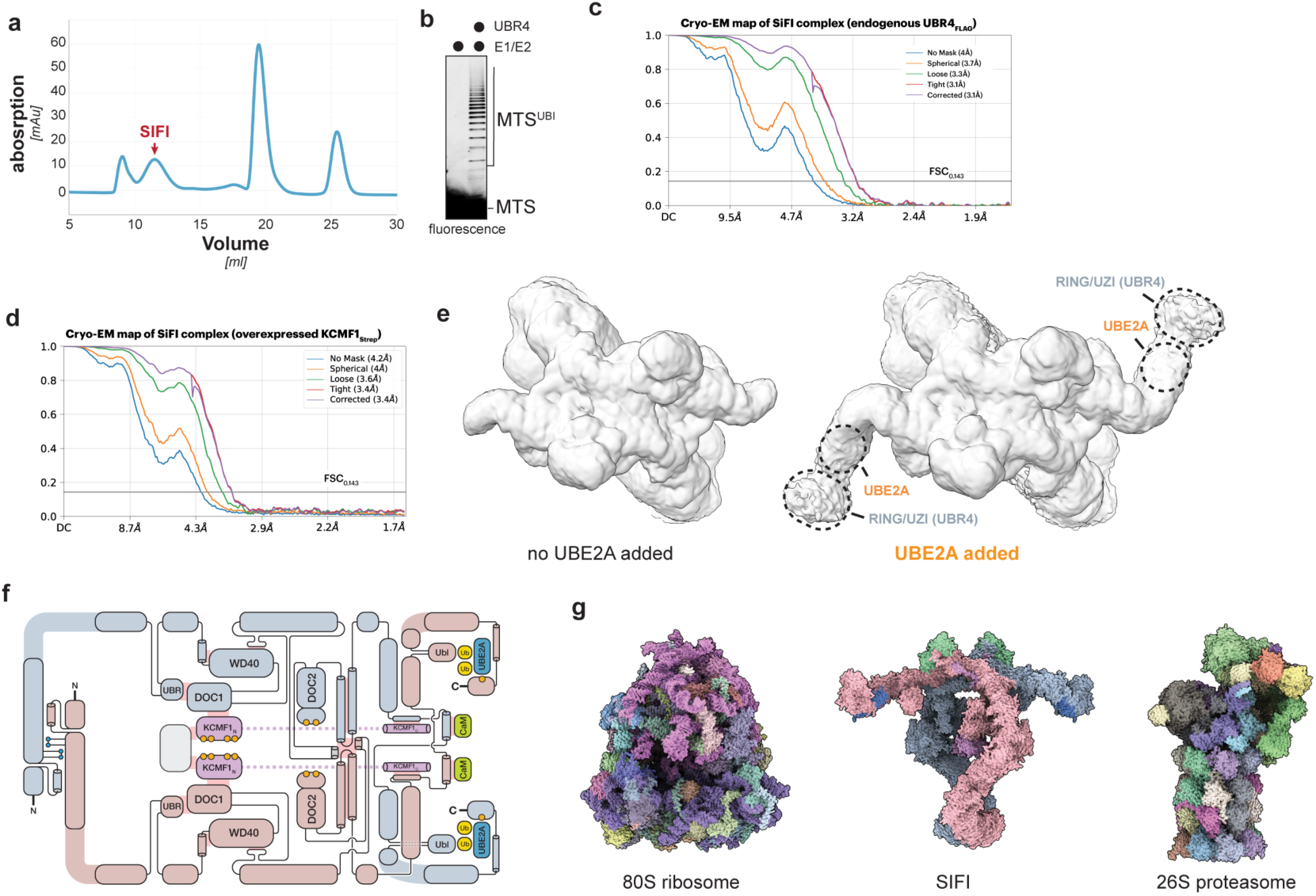
Structural characterization of human SIFI. **a.** Size exclusion chromatography of affinity-purified endogenous UBR4^FLAG^. Red arrow denotes the SIFI peak, confirmed by Western blotting and negative stain EM (data not shown). **b.** *In vitro* ubiquitylation of a fluorescently labeled presequence peptide shows that SEC purified endogenous SIFI used for cryo-EM is active in catalyzing ubiquitylation. Similar results in n=2 independent experiments. **c.** Fourier shell correlation (FSC) plot of the cryo-EM map of endogenous SIFI complexes. Resolution cutoff for gold-standard FSC0.143 is shown. **d.** FSC plot of the cryo-EM map of SIFI complexes purified by affinity-purification of overexpressed KCMF1^Strep^. **e.** Comparison of the cryo-EM maps of SIFI (left) and SIFI supplemented with recombinant UBE2A (right) reveals stabilization of the C-terminal region of UBR4, including hemi-RING/UZI domain. **f.** Schematic representation of the SIFI complex. **g.** Comparison between SIFI, 26S proteasome (PDB 6MSB), and 80S ribosome (PDB 6QZP) reveals their similar size dimensions.

**Extended Data Figure 2:**
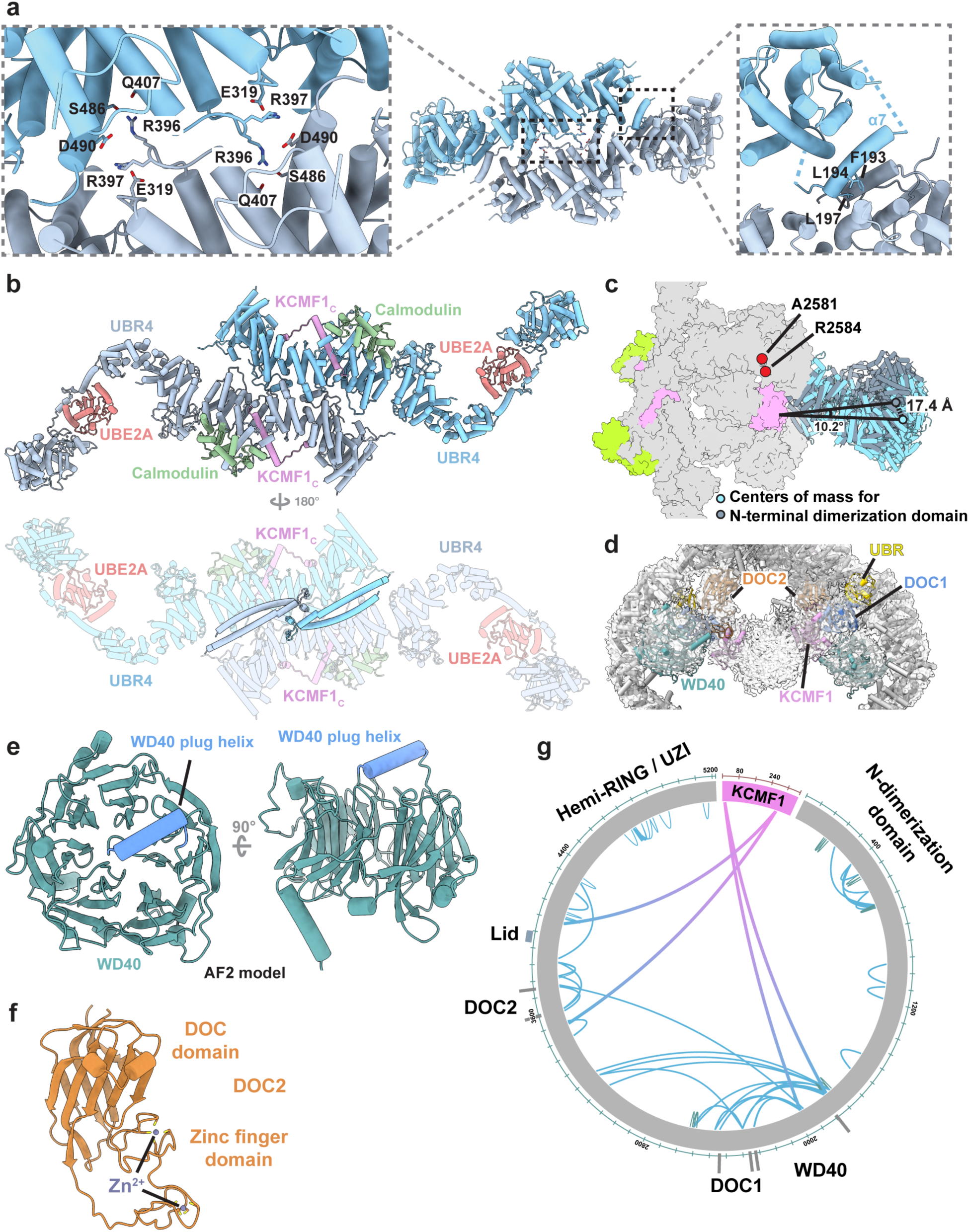
Structural domains of SIFI. **a.** Structural insights into the N-terminal dimerization region of SIFI. *Left*: two Lys residues at the tips of interlocking loops from each protomer engage in polar interactions with their paired counterpart. *Right*: α7 helix of UBR4 undergoes a helix swap with the paired protomer to form a helical bundle. **b.** C-terminal dimerization region. 180° rotation reveals the stabilizing helical pair at the inside of the SIFI scaffold. **c.** The SIFI scaffold shows flexibility around a central hinge region, leading to ∼10° rotations of the N- and C- terminal dimerization domains relative to each other. Two disease mutations in *UBR4*, A2581 (ataxia) and R2584 (cancer) are located close to the hinge region. **d.** Cryo-EM map of central SIFI showing docked WD40, DOC and UBR domains. These internal protein interaction modules are flexible and hence show lower overall resolution. The central density (ABHD10) is described below. **e.** The WD40 domain of UBR4 features a central plug helix at a surface commonly involved in protein interactions characteristic of WD40 repeats. **f.** The DOC2 domain of UBR4 contains two Zinc-binding loops that collectively form a potential protein interaction interface. **g.** Diagram of cross-linking mass spectrometry validates domain arrangement at the center of SIFI.

**Extended Data Figure 3:**
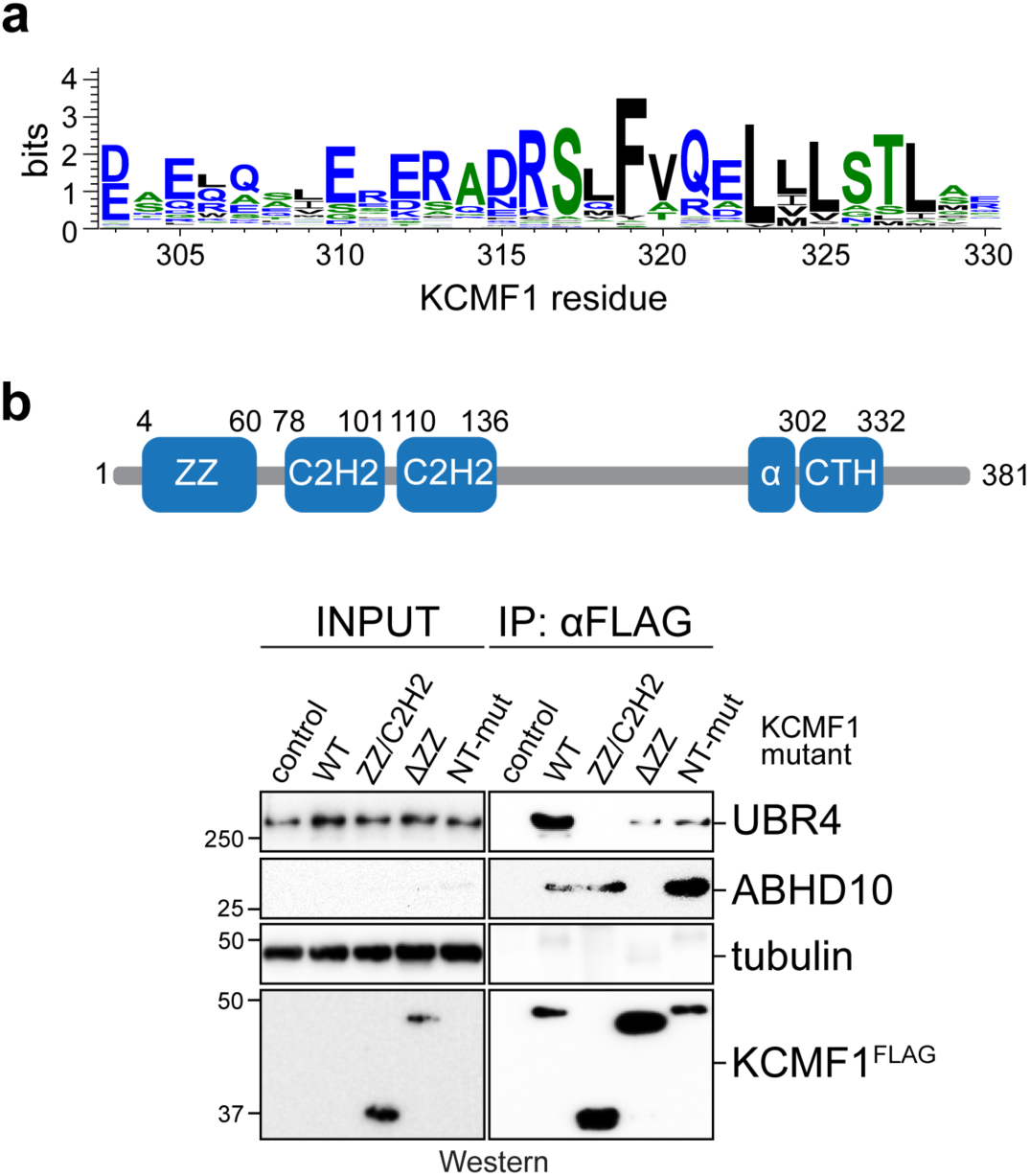
KCMF1 recruitment and function in SIFI. **a.** The central Phe residue in the C-terminal KCMF1 helix is highly conserved. **b.** *top:* Schematic of domain arrangement in KCMF1. *bottom:* Deletion of the N-terminal ZZ domain or mutation of R20E, E38R, L61E, Y64E, R68E in the ZZ domain strongly impaired the interaction between KCMF1^FLAG^ and endogenous UBR4, as shown by affinity-purification and Western blotting. The ZZ deletion also interfered with recognition of the N-degron substrate ABHD10, as described below. Similar results in n=2 independent experiments.

**Extended Data Figure 4:**
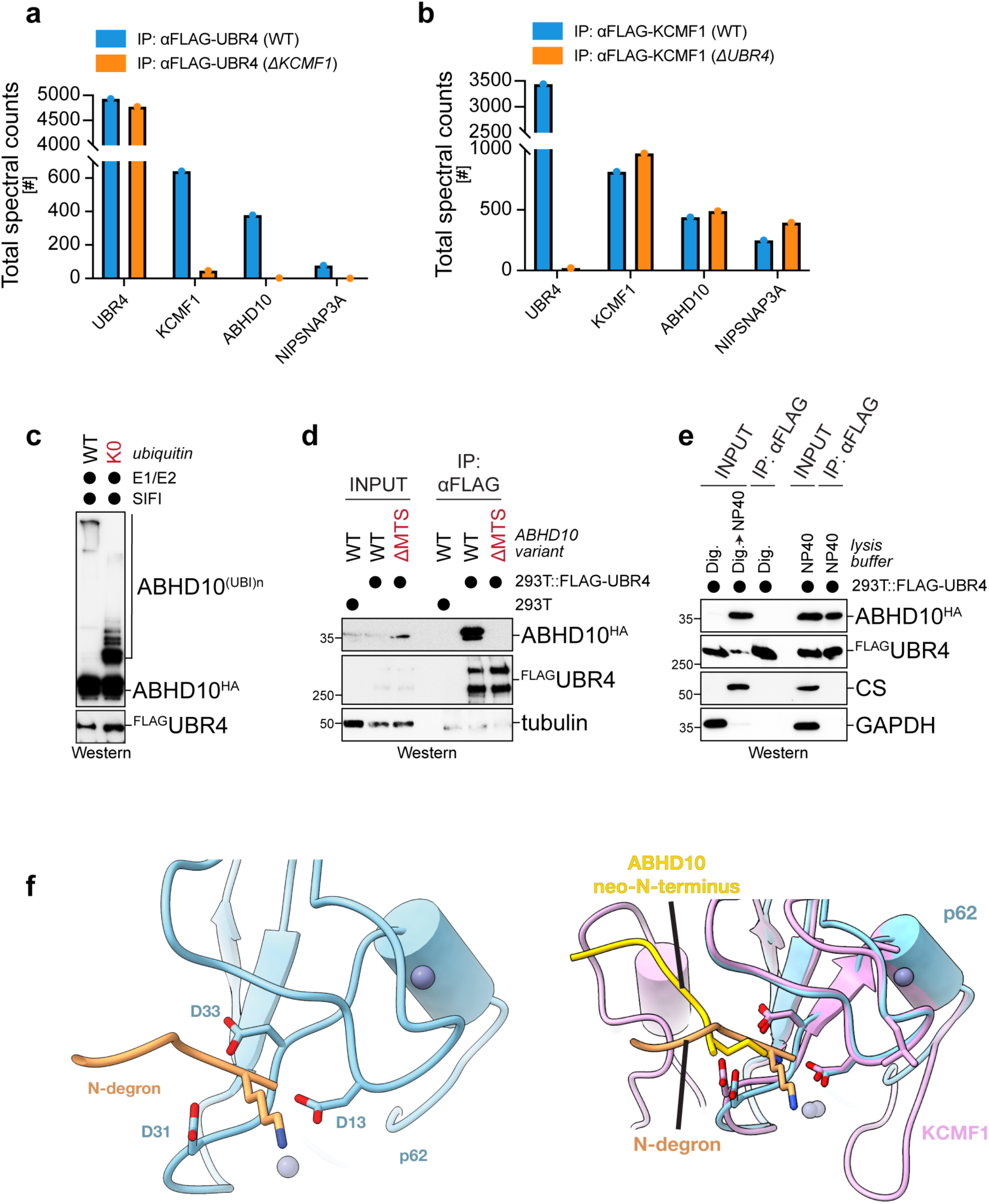
ABHD10 binds SIFI as an N-degron substrate. **a.** Immunoprecipitation of SIFI in WT or *ΔKCMF1* cells coupled to mass spectrometry shows that two mitochondrial proteins, ABHD10 and NIPSNAP3A, are lost from SIFI purifications in the absence of KCMF1. **b.** Immunoprecipitation of KCMF1 coupled to mass spectrometry from WT or *ΔUBR4* cells shows that KCMF1 binds to ABHD10 and NIPSNAP3A independently of UBR4. **c.** SIFI-bound ABHD10 is ubiquitylated if SIFI is incubated with E1, UBE2D3/UBE2A, and ubiquitin, as shown by Western blotting. Ubiquitin lacking Lys residues and hence unable to support chain formation (K0) is used as control. Similar results in n=2 independent experiments. **d.** ABHD10 only binds KCMF1 if it contains an N-terminal presequence that is cleaved off upon entry into mitochondria. ΔMTS-ABHD10 does not contain its presequence and is not imported into mitochondria, thus initiating with a Met residue. Binding of ABHD10^HA^ to KCMF1^FLAG^ was analyzed by αFLAG-affinity-purification and Western blotting. Similar results in n=2 independent experiments. **e.** ABHD10 only binds KCMF1 after being released from mitochondria. Cells were either lysed with digitonin, which preserves mitochondrial integrity, or NP40, which disrupts mitochondrial integrity and results in release of mitochondrial proteins into the lysate. UBR4^FLAG^ was affinity-purified and bound ABHD10^HA^ was detected by Western blotting. Similar results in n=2 independent experiments. **f.** *Left:* Structure of the ZZ-domain of p62, which is known to bind to N-degrons as determined by X-ray crystallography. *Right:* overlay of p62 and KCMF1 N-domains bound to their respective N-degrons.

**Extended Data Figure 5:**
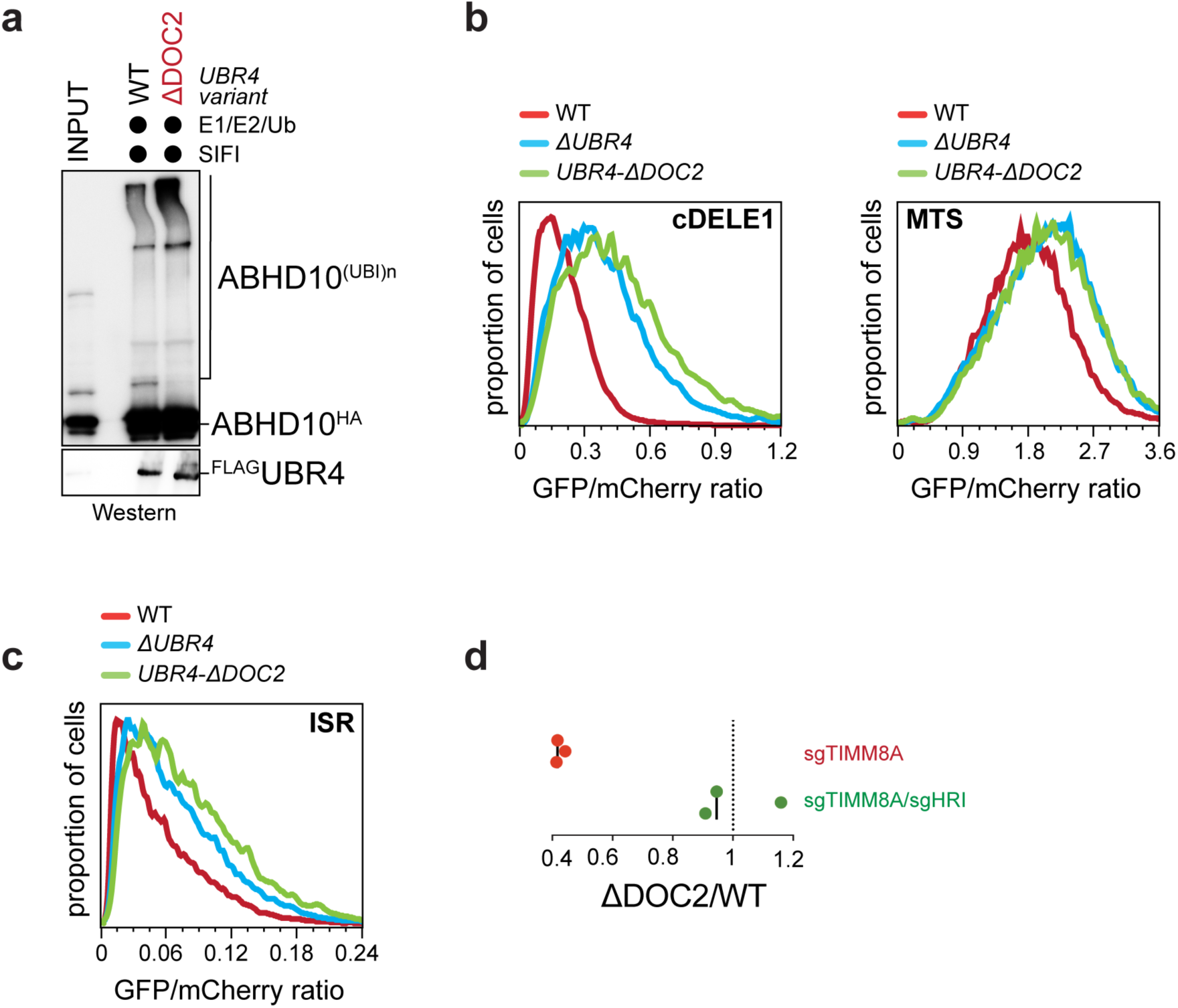
The DOC2 domain of SIFI is important for HRI ubiquitylation and degradation. **a.** The DOC2 domain of SIFI is not required for ABHD10 ubiquitylation. SIFI was purified from either WT or *UBR4*^ΔDOC2^ cells that also expressed ABHD10^HA^ and incubated with E1, UBE2D3/UBE2A, and ubiquitin. Ubiquitylation of bound ABDH10 was detected by Western blotting. Experiment performed once. **b.** The DOC2 domain of SIFI is required for degradation of DELE1 or unimported mito-chondrial proteins. The stability of DELE1 or the presequence of COX8A was determined in WT or *UBR4*^ΔDOC2^ cells by flow cytometry, as described before. Similar results in n=2 independent experiments. **c.** Deletion of the DOC2 domain in *UBR4* results in stronger activation of the integrated stress response. A reporter for integrated stress response activation based on uORFs of ATF4 was expressed in either WT, *ΔUBR4*, or *UBR4*^ΔDOC2^ cells, which were treated with sodium arsenite and analyzed by flow cytometry. Similar results in n=2 independent experiments. **d.** The DOC2 domain of UBR4 is essential for cell survival upon mitochondrial import stress. WT and *UBR4*^ΔDOC2^ cells were labeled with GFP and mCherry, respectively, mixed, and depleted of either TIMM8A or TIMM8A/HRI, as indicated. After 12d of co-culture, the ratio between cell types was determined by flow cytometry. Each datapoint represents a biological replicate.

**Extended Data Figure 6:**
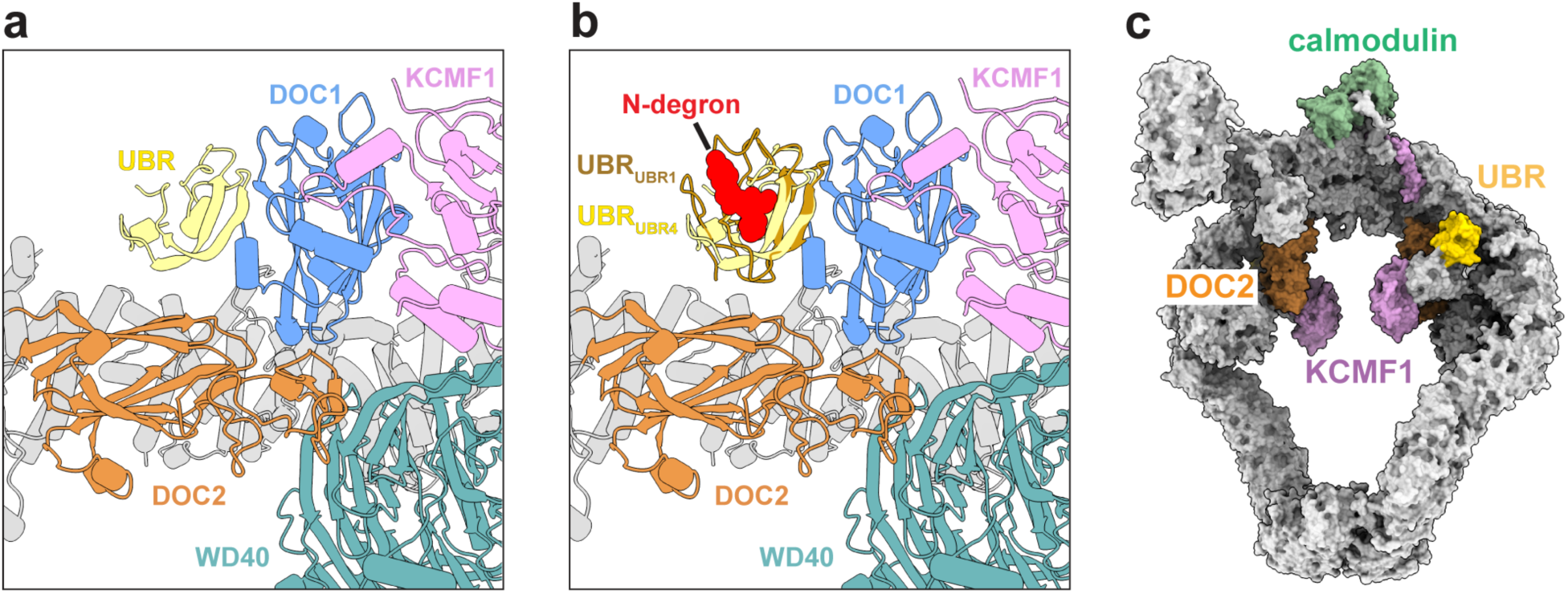
The UBR boxes of UBR4 present substrates towards the interior of SIFI. **a.** The UBR box of UBR4 binds to DOC1 domain of UBR4 and the N-terminal domain of KCMF1, forming the KCMF1^N138^-DOC1-UBR subcomplex. X-ray structure of the UBR box of UBR1 (PDB 3NIH)^32^ reveals that its peptide binding groove is open. **b.** Modeling a N-degron peptide into the open substrate-binding groove of the UBR box of UBR4, based on the UBR1 X-ray structure (PDB 3NIH). **c.** Structure of SIFI showing the proximity of UBR box to other substrate-binding modules of SIFI, including the DOC2 domain of UBR4 and the ZZ domain of KCMF1.

**Extended Data Figure 7:**
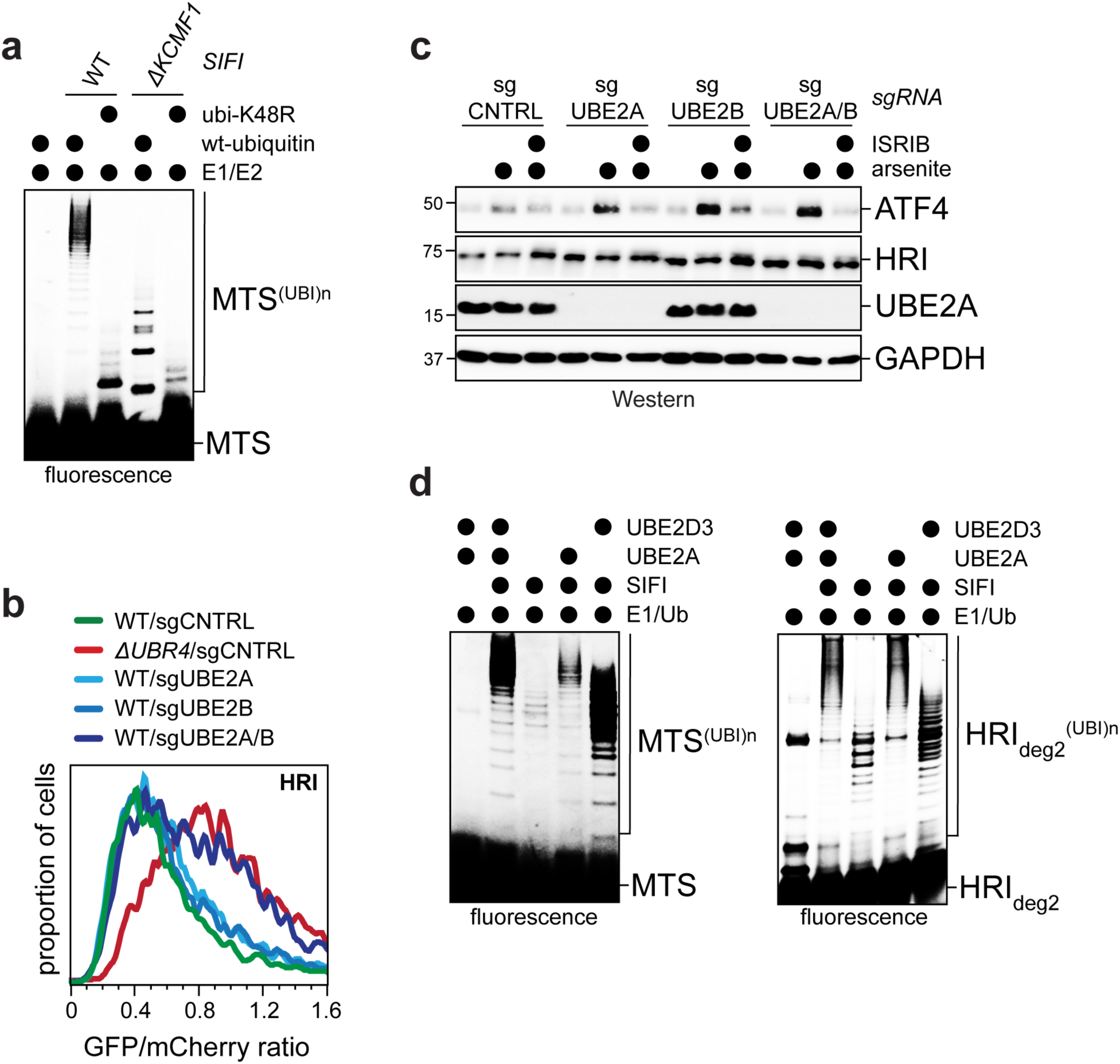
UBE2A promotes ubiquitin chain elongation by SIFI. **a.** KCMF1 does not determine linkage specificity of SIFI. SIFI was purified from WT or *ΔKCMF1* cells and tested for its ability to catalyze ubiquitylation of a mitochondrial presequence. As indicated, either wt-ubiquitin or ubiquitin-K48R were used. **b.** UBE2A and its close homolog UBE2B are required for HRI degradation. UBE2A and UBE2B were co-depleted from 293T cells, and HRI stability was monitored using a GFP-HRI::mCherry reporter, as described before. Similar results in n=2 independent experiments. **c.** UBE2A and UBE2B co-depletion results in increased activation of the integrated stress response, as determined by αATF4-Western blotting. Experiment performed once. **d.** UBE2A promotes ubiquitin chain elongation. SIFI was affinity-purified via UBR4^FLAG^ and incubated with E1, ubiquitin and a mitochondrial presequence peptide (left) or HRI degron helix 2 peptide (right), resulting in short ubiquitin chains on the presequence substrate likely catalyzed by an E2 that co-purified with SIFI. Addition of the E2 UBE2D3 results in an increase in short ubiquitin chains, indicative of a role of UBE2D3 in chain initiation. Addition of UBE2A leads to selective elongation of ubiquitin chains that had already been initiated. Similar results in n=2 independent experiments.

**Extended Data Figure 8:**
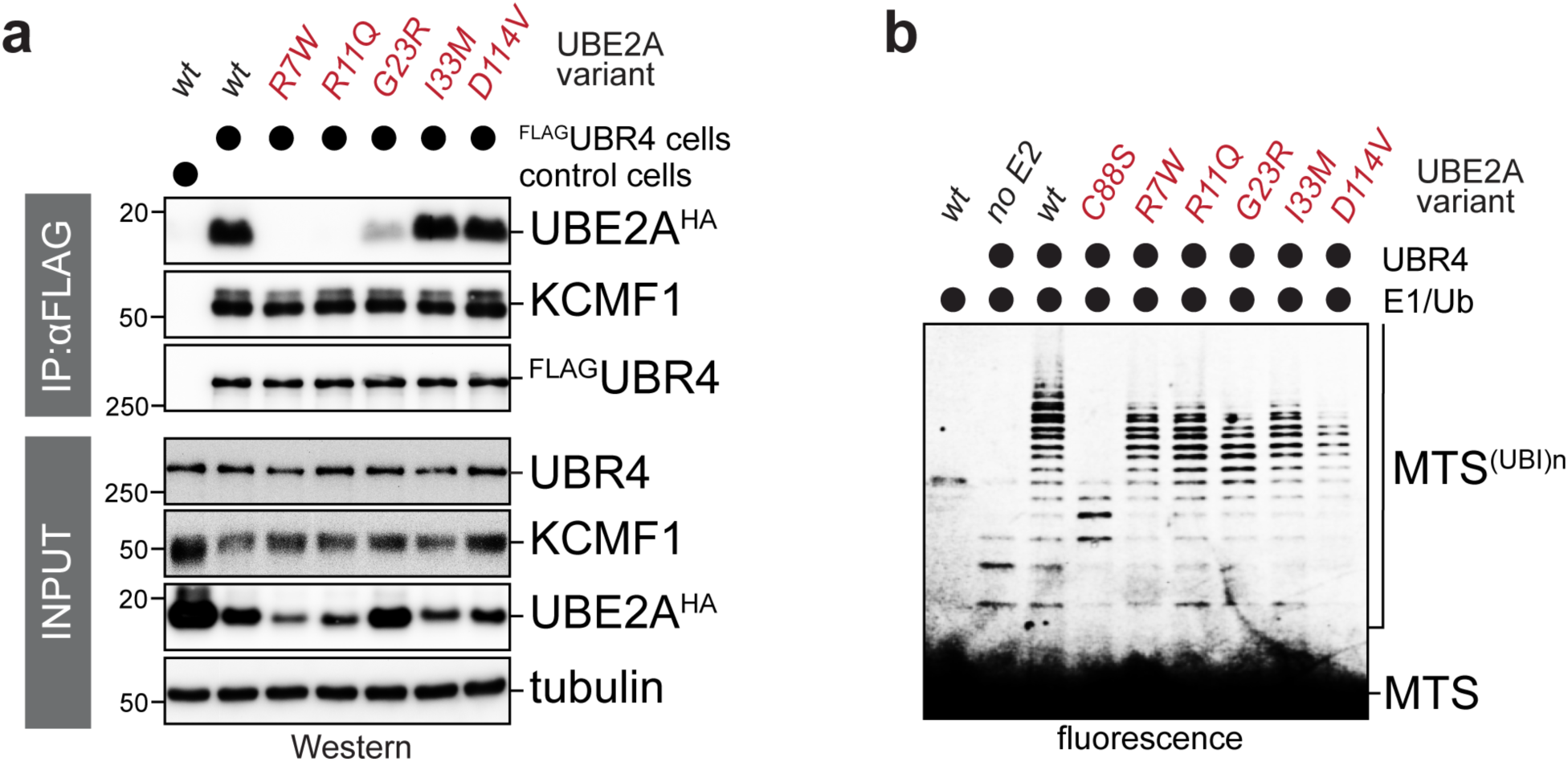
Validation of the UBE2A embrace by UBR4. **a.** UBE2A mutants detected in patients of Nascimento Syndrome are impaired in binding SIFI in cells, as determined by UBR4 affinity-purification and Western blotting. Similar results in n=2 independent experiments. **b.** UBE2A variants in Nascimento Syndrome are impaired, but not fully inhibited, in catalyzing ubiquitin chain elongation on SIFI substrates. D114 of UBE2A is at the interface with donor ubiquitin. Similar results in n=2 independent experiments.

**Extended Data Figure 9:**
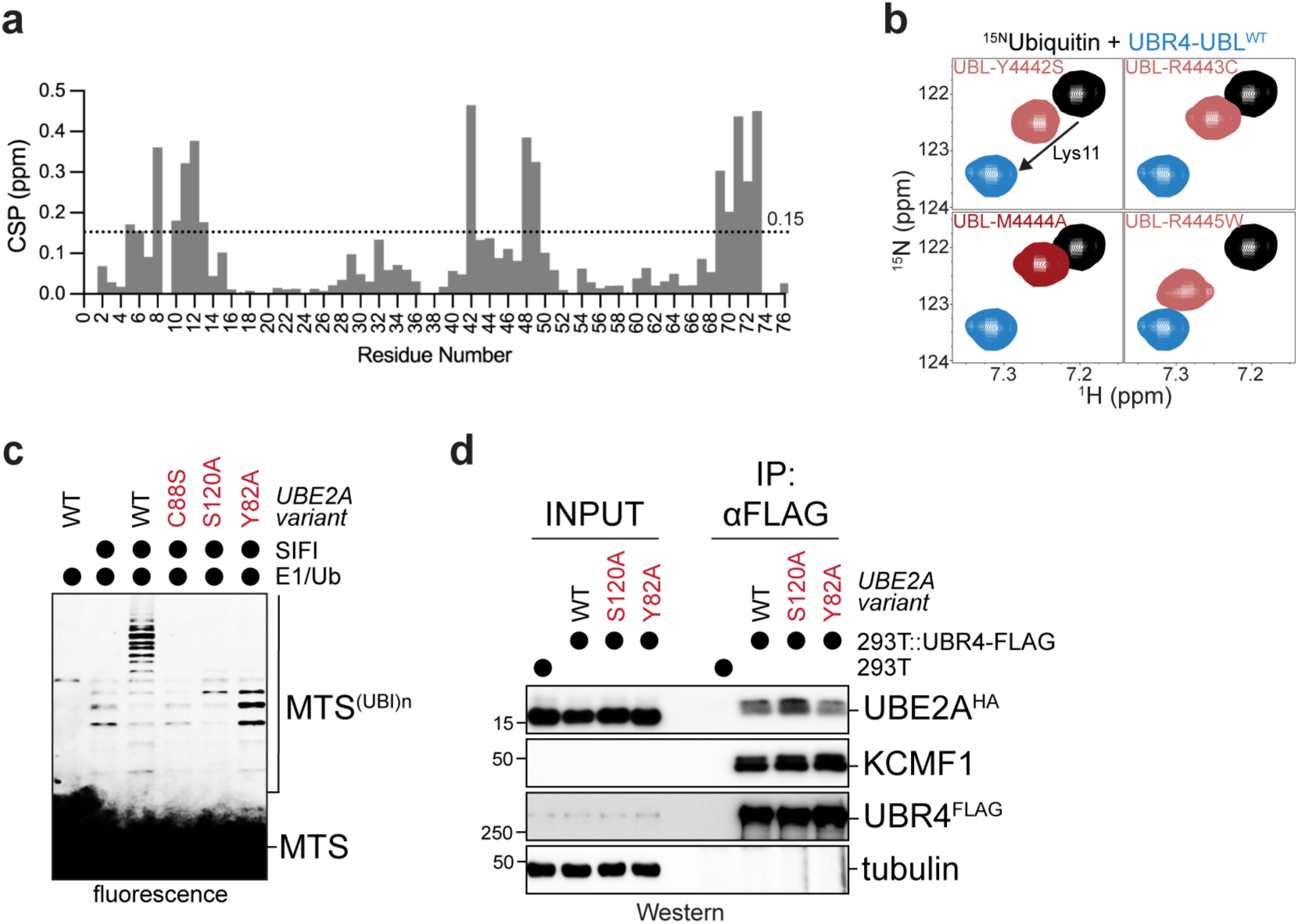
The UBL domain of UBR4 orients an acceptor ubiquitin-K48 to the active site of UBE2A. **a.** Chemical shift perturbation analysis of ^15^N-ubiquitin with 2.5 molar equivalents of UBR4 UBL domain. **b.** NMR analysis of UBR4 UBL mutants binding to ^15^N-ubiquitin. Lys11 of ubiquitin was chosen as a reporter of binding because it is far away from the interface directly affected by UBL mutations. **c.** Y82 and S120 of UBE2A are essential for ubiquitin chain elongation by SIFI, as detected in an *in vitro* ubiquitylation assay as described above. Experiment performed once. **d.** Mutation of Y82 or S120 of UBE2A does not prevent binding of the E2 to SIFI. Endogenous SIFI was purified from UBR4^FLAG^ cells and co-purifying UBE2A^HA^ variants were detected by Western blotting. Experiment performed once.

**Extended Data Fig. 10:**
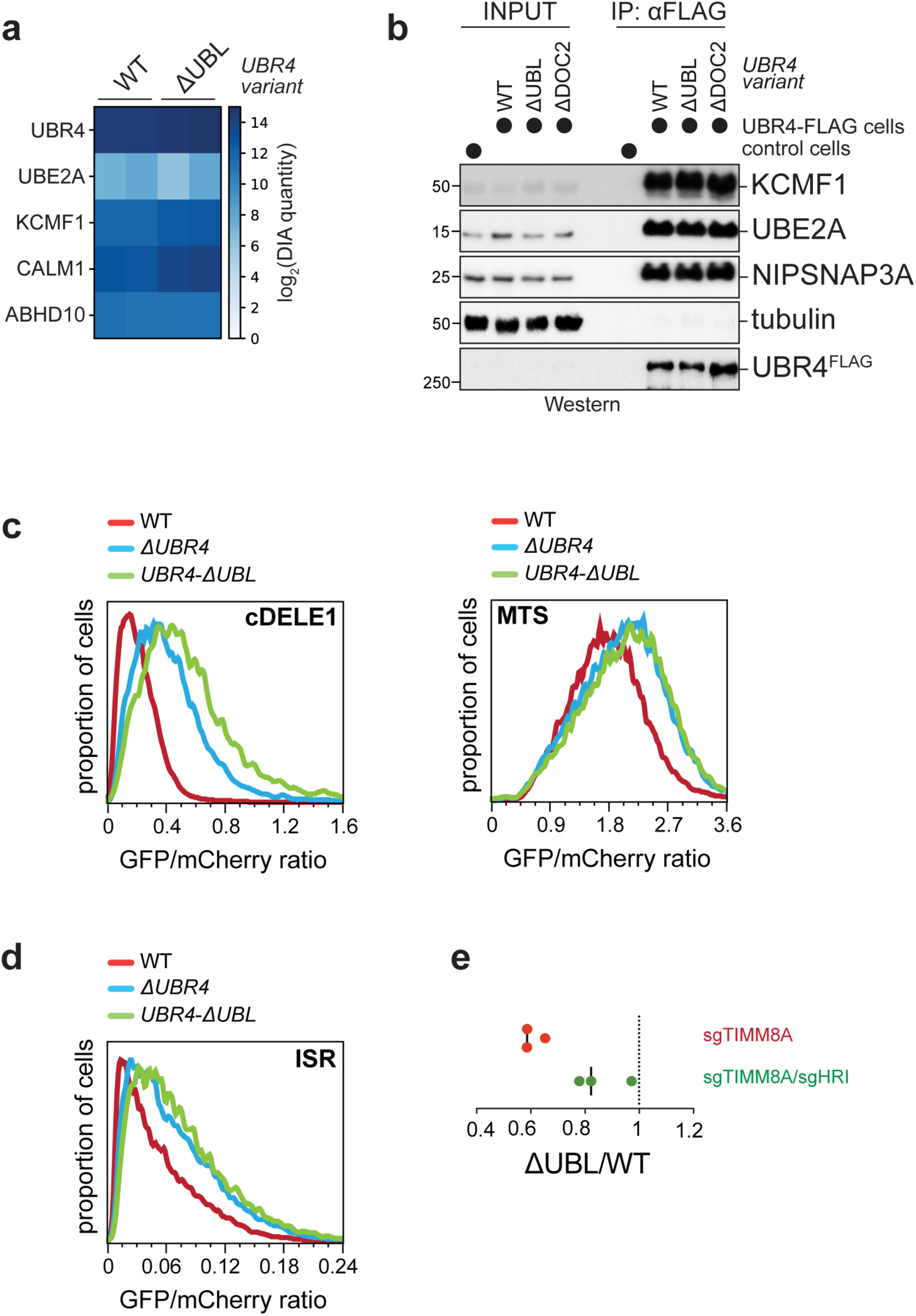
The UBL domain of UBR4 is essential for ubiquitin chain elongation. **a.** Mass spectrometry of SIFI^ΔUBL^ indicates that the UBL domain is not required for structural integrity of SIFI. Endogenous SIFI was affinity-purified from WT or *UBR4^ΔUBL^* cells and bound proteins were detected by mass spectrometry. Technical replicates are shown. **b.** Western blotting of SIFI^ΔUBL^ shows that the UBL domain is not required for structural integrity of SIFI. Endogenous SIFI was affinity-purified from WT or *UBR4^ΔUBL^* cells and bound proteins were detected by Western blotting. Experiment performed once and validated by mass spectrometry. **c.** The UBL domain of UBR4 is required for degradation of DELE1 or unimported mitochondrial proteins. Protein stability was determined by flow cytometry, as described before. Similar results in n=2 independent experiments. **d.** Deletion of the UBL domain in UBR4 increases stress response activation upon sodium arsenite treatment, as detected by uORF-ATF4 reporters in flow cytometry as described above. Similar results in n=2 independent experiments. **e.** The UBL domain of UBR4 is essential for cell survival upon mitochondrial import stress. WT and *UBR4*^ΔUBL^ cells were labeled with GFP and mCherry, respectively, mixed, and depleted of either TIMM8A or TIMM8A/HRI, as indicated. After 12d of co-culture, the ratio between cell types was determined by flow cytometry. Each datapoint represents a biological replicate.

## References

1. Costa-Ma+oli, M., and Walter, P. (2020). 5314. Science 368. 10.1126/science.aat5314.

2. Harper, J.W., and Bennett, E.J. (2016). Proteome complexity and the forces that drive proteome imbalance. Nature 537, 328–338. 10.1038/nature19947.

3. Hipp, M.S., Kasturi, P., and Hartl, F.U. (2019). The proteostasis network and its decline in ageing. Nat Rev Mol Cell Biol 20, 421–435. 10.1038/s41580-019-0101-y.

4. Fessler, E., Eckl, E.M., Schmitt, S., Mancilla, I.A., Meyer-Bender, M.F., Hanf, M., Philippou-Massier, J., Krebs, S., Zischka, H., and Jae, L.T. (2020). A pathway coordinated by DELE1 relays mitochondrial stress to the cytosol. Nature 579, 433–437. 10.1038/s41586-020-2076-4.

5. Guo, X., Aviles, G., Liu, Y., Tian, R., Unger, B.A., Lin, Y.T., Wiita, A.P., Xu, K., Correia, M.A., and Kampmann, M. (2020). Mitochondrial stress is relayed to the cytosol by an OMA1-DELE1-HRI pathway. Nature 579, 427–432. 10.1038/s41586-020-2078-2.

6. McEwen, E., Kedersha, N., Song, B., Scheuner, D., Gilks, N., Han, A., Chen, J.J., Anderson, P., and Kaufman, R.J. (2005). Heme-regulated inhibitor kinase-mediated phosphorylation of eukaryotic translation initiation factor 2 inhibits translation, induces stress granule formation, and mediates survival upon arsenite exposure. J Biol Chem 280, 16925–16933. 10.1074/jbc.M412882200.

7. Anderson, N.S., and Haynes, C.M. (2020). Folding the Mitochondrial UPR into the Integrated Stress Response. Trends Cell Biol 30, 428–439. 10.1016/j.tcb.2020.03.001.

8. Winter, J.M., Yadav, T., and Rutter, J. (2022). Stressed to death: Mitochondrial stress responses connect respiration and apoptosis in cancer. Mol Cell 82, 3321–3332. 10.1016/j.molcel.2022.07.012.

9. Inada, T., and Beckmann, R. (2024). Mechanisms of Translation-coupled Quality Control. J Mol Biol 436, 168496. 10.1016/j.jmb.2024.168496.

10. Haakonsen, D.L., Heider, M., Ingersoll, A.J., Vodehnal, K., Witus, S.R., Uenaka, T., Wernig, M., and Rape, M. (2024). Stress response silencing by an E3 ligase mutated in neurodegeneration. Nature 626, 874–880. 10.1038/s41586-023-06985-7.

11. Nakaya, T., Ishiguro, K., Belzil, C., Rietsch, A.M., Yu, Q., Mizuno, S., Bronson, R.T., Geng, Y., Nguyen, M.D., Akashi, K., et al. (2013). p600 Plays Essential Roles in Fetal Development. PLoS One 8, e66269. 10.1371/journal.pone.0066269.

12. Tasaki, T., Kim, S.T., Zakrzewska, A., Lee, B.E., Kang, M.J., Yoo, Y.D., Cha-Molstad, H.J., Hwang, J., Soung, N.K., Sung, K.S., et al. (2013). UBR box N-recognin-4 (UBR4), an N-recognin of the N-end rule pathway, and its role in yolk sac vascular development and autophagy. Proc Natl Acad Sci U S A 110, 3800–3805. 10.1073/pnas.1217358110.

13. Hunt, L.C., Stover, J., Haugen, B., Shaw, T.I., Li, Y., Pagala, V.R., Finkelstein, D., Barton, E.R., Fan, Y., Labelle, M., et al. (2019). A Key Role for the Ubiquitin Ligase UBR4 in Myofiber Hypertrophy in Drosophila and Mice. Cell Rep 28, 1268–1281 e1266. 10.1016/j.celrep.2019.06.094.

14. Hunt, L.C., Schadeberg, B., Stover, J., Haugen, B., Pagala, V., Wang, Y.D., Puglise, J., Barton, E.R., Peng, J., and Demontis, F. (2021). Antagonistic control of myofiber size and muscle protein quality control by the ubiquitin ligase UBR4 during aging. Nat Commun 12, 1418. 10.1038/s41467-021-21738-8.

15. Conroy, J., McGe+gan, P., Murphy, R., Webb, D., Murphy, S.M., McCoy, B., Albertyn, C., McCreary, D., McDonagh, C., Walsh, O., et al. (2014). A novel locus for episodic ataxia:UBR4 the likely candidate. Eur J Hum Genet 22, 505–510. 10.1038/ejhg.2013.173.

16. Yau, R.G., Doerner, K., Castellanos, E.R., Haakonsen, D.L., Werner, A., Wang, N., Yang, X.W., Martinez-Martin, N., Matsumoto, M.L., Dixit, V.M., and Rape, M. (2017). Assembly and Function of Heterotypic Ubiquitin Chains in Cell-Cycle and Protein Quality Control. Cell 171, 918–933 e920. 10.1016/j.cell.2017.09.040.

17. Abdel-Nour, M., Carneiro, L.A.M., Downey, J., Tsalikis, J., Outlioua, A., Prescott, D., Da Costa, L.S., Hovingh, E.S., Farahvash, A., Gaudet, R.G., et al. (2019). The heme-regulated inhibitor is a cytosolic sensor of protein misfolding that controls innate immune signaling. Science 365. 10.1126/science.aaw4144.

18. Girardin, S.E., Cuziol, C., Philpott, D.J., and Arnoult, D. (2021). The eIF2alpha kinase HRI in innate immunity, proteostasis, and mitochondrial stress. FEBS J 288, 3094–3107. 10.1111/febs.15553.

19. Yamano, K., and Youle, R.J. (2013). PINK1 is degraded through the N-end rule pathway. Autophagy 9, 1758–1769. 10.4161/auto.24633.

20. Barnsby-Greer, L., Mabbitt, P.D., Dery, M.A., Squair, D.R., Wood, N.T., Lamoliatte, F., Lange, S.M., and Virdee, S. (2024). UBE2A and UBE2B are recruited by an atypical E3 ligase module in UBR4. Nat Struct Mol Biol 31, 351–363. 10.1038/s41594-023-01192-4.

21. Tate, J.G., Bamford, S., Jubb, H.C., Sondka, Z., Beare, D.M., Bindal, N., Boutselakis, H., Cole, C.G., Creatore, C., Dawson, E., et al. (2019). COSMIC: the Catalogue Of Somatic Mutations In Cancer. Nucleic Acids Res 47, D941–D947. 10.1093/nar/gky1015.

22. Choi, K.D., Kim, J.S., Kim, H.J., Jung, I., Jeong, S.H., Lee, S.H., Kim, D.U., Kim, S.H., Choi, S.Y., Shin, J.H., et al. (2017). Genetic Variants Associated with Episodic Ataxia in Korea. Sci Rep 7, 13855. 10.1038/s41598-017-14254-7.

23. Meyers, R.M., Bryan, J.G., McFarland, J.M., Weir, B.A., Sizemore, A.E., Xu, H., Dharia, N.V., Montgomery, P.G., Cowley, G.S., Pantel, S., et al. (2017). Computational correction of copy number effect improves specificity of CRISPR-Cas9 essentiality screens in cancer cells. Nat Genet 49, 1779–1784. 10.1038/ng.3984.

24. Villalobo, A., Ishida, H., Vogel, H.J., and Berchtold, M.W. (2018). Calmodulin as a protein linker and a regulator of adaptor/scaffold proteins. Biochim Biophys Acta Mol Cell Res 1865, 507–521. 10.1016/j.bbamcr.2017.12.004.

25. Bhattacharyya, M., Karandur, D., and Kuriyan, J. (2020). Structural Insights into the Regulation of Ca(2+)/Calmodulin-Dependent Protein Kinase II (CaMKII). Cold Spring Harb Perspect Biol 12. 10.1101/cshperspect.a035147.

26. Belzil, C., Neumayer, G., Vassilev, A.P., Yap, K.L., Konishi, H., Rivest, S., Sanada, K., Ikura, M., Nakatani, Y., and Nguyen, M.D. (2013). A Ca2+-dependent mechanism of neuronal survival mediated by the microtubule-associated protein p600. J Biol Chem 288, 24452–24464. 10.1074/jbc.M113.483107.

27. Cha-Molstad, H., Yu, J.E., Feng, Z., Lee, S.H., Kim, J.G., Yang, P., Han, B., Sung, K.W., Yoo, Y.D., Hwang, J., et al. (2017). p62/SQSTM1/Sequestosome-1 is an N-recognin of the N-end rule pathway which modulates autophagosome biogenesis. Nat Commun 8, 102. 10.1038/s41467-017-00085-7.

28. Pierce, N.W., Kleiger, G., Shan, S.O., and Deshaies, R.J. (2009). Detection of sequential polyubiquitylation on a millisecond timescale. Nature 462, 615–619. nature08595 [pii] 10.1038/nature08595.

29. Jeong, D.E., Lee, H.S., Ku, B., Kim, C.H., Kim, S.J., and Shin, H.C. (2023). Insights into the recognition mechanism in the UBR box of UBR4 for its specific substrates. Commun Biol 6, 1214. 10.1038/s42003-023-05602-7.

30. Tasaki, T., Mulder, L.C., Iwamatsu, A., Lee, M.J., Davydov, I.V., Varshavsky, A., Muesing, M., and Kwon, Y.T. (2005). A family of mammalian E3 ubiquitin ligases that contain the UBR box motif and recognize N-degrons. Mol Cell Biol 25, 7120–7136. 10.1128/MCB.25.16.7120-7136.2005.

31. Heo, A.J., Kim, S.B., Ji, C.H., Han, D., Lee, S.J., Lee, S.H., Lee, M.J., Lee, J.S., Ciechanover, A., Kim, B.Y., and Kwon, Y.T. (2021). The N-terminal cysteine is a dual sensor of oxygen and oxidative stress. Proc Natl Acad Sci U S A 118. 10.1073/pnas.2107993118.

32. Choi, W.S., Jeong, B.C., Joo, Y.J., Lee, M.R., Kim, J., Eck, M.J., and Song, H.K. (2010). Structural basis for the recognition of N-end rule substrates by the UBR box of ubiquitin ligases. Nat Struct Mol Biol 17, 1175–1181. 10.1038/nsmb.1907.

33. Budny, B., Badura-Stronka, M., Materna-Kiryluk, A., Tzschach, A., Raynaud, M., Latos-Bielenska, A., and Ropers, H.H. (2010). Novel missense mutations in the ubiquitination-related gene UBE2A cause a recognizable X-linked mental retardation syndrome. Clin Genet 77, 541–551. 10.1111/j.1399-0004.2010.01429.x.

34. Cordeddu, V., Macke, E.L., Radio, F.C., Lo Cicero, S., Pantaleoni, F., Ta+, M., Bellacchio, E., Ciolfi, A., Agolini, E., Bruselles, A., et al. (2020). Refinement of the clinical and mutational spectrum of UBE2A deficiency syndrome. Clin Genet 98, 172–178. 10.1111/cge.13775.

35. Nascimento, R.M., Otto, P.A., de Brouwer, A.P., and Vianna-Morgante, A.M. (2006). UBE2A, which encodes a ubiquitin-conjugating enzyme, is mutated in a novel X-linked mental retardation syndrome. American journal of human genetics 79, 549–555. 10.1086/507047.

36. Deng, Z., Ai, H., Sun, M., Tong, Z., Du, Y., Qu, Q., Zhang, L., Xu, Z., Tao, S., Shi, Q., et al. (2023). Mechanistic insights into nucleosomal H2B monoubiquitylation mediated by yeast Bre1-Rad6 and its human homolog RNF20/RNF40-hRAD6A. Mol Cell 83, 3080–3094 e3014. 10.1016/j.molcel.2023.08.001.

37. Hibbert, R.G., Huang, A., Boelens, R., and Sixma, T.K. (2011). E3 ligase Rad18 promotes monoubiquitination rather than ubiquitin chain formation by E2 enzyme Rad6. Proc Natl Acad Sci U S A 108, 5590–5595. 1017516108 [pii] 10.1073/pnas.1017516108.

38. Kumar, B., Lecompte, K.G., Klein, J.M., and Haas, A.L. (2010). Ser(120) of Ubc2/Rad6 regulates ubiquitin-dependent N-end rule targeting by E3alpha/Ubr1. J Biol Chem 285, 41300–41309. 10.1074/jbc.M110.169136.

39. Leto, D.E., Morgens, D.W., Zhang, L., Walczak, C.P., Elias, J.E., Bassik, M.C., and Kopito, R.R. (2019). Genome-wide CRISPR Analysis Identifies Substrate-Specific Conjugation Modules in ER-Associated Degradation. Mol Cell 73, 377–389 e311. 10.1016/j.molcel.2018.11.015.

40. Eletr, Z.M., Huang, D.T., Duda, D.M., Schulman, B.A., and Kuhlman, B. (2005). E2 conjugating enzymes must disengage from their E1 enzymes before E3-dependent ubiquitin and ubiquitin-like transfer. Nature structural & molecular biology 12, 933–934. 10.1038/nsmb984.

41. Kamadurai, H.B., Souphron, J., Scott, D.C., Duda, D.M., Miller, D.J., Stringer, D., Piper, R.C., and Schulman, B.A. (2009). Insights into ubiquitin transfer cascades from a structure of a UbcH5B approximately ubiquitin-HECT(NEDD4L) complex. Mol Cell 36, 1095–1102. S1097-2765(09)00824-7 [pii] 10.1016/j.molcel.2009.11.010.

42. Koegl, M., Hoppe, T., Schlenker, S., Ulrich, H.D., Mayer, T.U., and Jentsch, S. (1999). A novel ubiquitination factor, E4, is involved in multiubiquitin chain assembly. Cell 96, 635–644. S0092-8674(00)80574-7 [pii].

43. Kaiho-Soma, A., Akizuki, Y., Igarashi, K., Endo, A., Shoda, T., Kawase, Y., Demizu, Y., Naito, M., Saeki, Y., Tanaka, K., and Ohtake, F. (2021). TRIP12 promotes small-molecule-induced degradation through K29/K48-branched ubiquitin chains. Mol Cell 81, 1411–1424 e1417. 10.1016/j.molcel.2021.01.023.

44. Wickliffe, K.E., Lorenz, S., Wemmer, D.E., Kuriyan, J., and Rape, M. (2011). The mechanism of linkage-specific ubiquitin chain elongation by a single-subunit e2. Cell 144, 769–781. S0092-8674(11)00111-5 [pii] 10.1016/j.cell.2011.01.035.

45. Eddins, M.J., Carlile, C.M., Gomez, K.M., Pickart, C.M., and Wolberger, C. (2006). Mms2-Ubc13 covalently bound to ubiquitin reveals the structural basis of linkage-specific polyubiquitin chain formation. Nat Struct Mol Biol 13, 915–920. nsmb1148 [pii] 10.1038/nsmb1148.

46. Hodakova, Z., Grishkovskaya, I., Brunner, H.L., Bolhuis, D.L., Belacic, K., Schleiffer, A., Kotisch, H., Brown, N.G., and Haselbach, D. (2023). Cryo-EM structure of the chain-elongating E3 ubiquitin ligase UBR5. EMBO J 42, e113348. 10.15252/embj.2022113348.

47. Hehl, L.A., Horn-Ghetko, D., Prabu, J.R., Vollrath, R., Vu, D.T., Perez Berrocal, D.A., Mulder, M.P.C., van der Heden van Noort, G.J., and Schulman, B.A. (2024). Structural snapshots along K48-linked ubiquitin chain formation by the HECT E3 UBR5. Nat Chem Biol 20, 190–200. 10.1038/s41589-023-01414-2.

48. Mark, K.G., Kolla, S., Aguirre, J.D., Garshott, D.M., Schmitt, S., Haakonsen, D.L., Xu, C., Kater, L., Kempf, G., Martinez-Gonzalez, B., et al. (2023). Orphan quality control shapes network dynamics and gene expression. Cell 186, 3460–3475 e3423. 10.1016/j.cell.2023.06.015.

49. Liu, C., Liu, W., Ye, Y., and Li, W. (2017). Ufd2p synthesizes branched ubiquitin chains to promote the degradation of substrates modified with atypical chains. Nat Commun 8, 14274. 10.1038/ncomms14274.

50. Kolla, S., Ye, M., Mark, K.G., and Rape, M. (2022). Assembly and function of branched ubiquitin chains. Trends Biochem Sci 47, 759–771. 10.1016/j.tibs.2022.04.003.

51. Jevtic, P., Haakonsen, D.L., and Rape, M. (2021). An E3 ligase guide to the galaxy of small-molecule-induced protein degradation. Cell Chem Biol. 10.1016/j.chembiol.2021.04.002.

52. Mastronarde, D.N. (2005). Automated electron microscope tomography using robust prediction of specimen movements. J Struct Biol 152, 36–51. 10.1016/j.jsb.2005.07.007.

53. Punjani, A., Rubinstein, J.L., Fleet, D.J., and Brubaker, M.A. (2017). cryoSPARC: algorithms for rapid unsupervised cryo-EM structure determination. Nat Methods 14, 290–296. 10.1038/nmeth.4169.

54. Punjani, A., Zhang, H., and Fleet, D.J. (2020). Non-uniform refinement: adaptive regularization improves single-particle cryo-EM reconstruction. Nat Methods 17, 1214–1221. 10.1038/s41592-020-00990-8.

55. Punjani, A., and Fleet, D.J. (2023). 3DFlex: determining structure and motion of flexible proteins from cryo-EM. Nat Methods 20, 860–870. 10.1038/s41592-023-01853-8.

56. Jumper, J., Evans, R., Pritzel, A., Green, T., Figurnov, M., Ronneberger, O., Tunyasuvunakool, K., Bates, R., Zidek, A., Potapenko, A., et al. (2021). Highly accurate protein structure prediction with AlphaFold. Nature 596, 583–589. 10.1038/s41586-021-03819-2.

57. Afonine, P.V., Poon, B.K., Read, R.J., Sobolev, O.V., Terwilliger, T.C., Urzhumtsev, A., and Adams, P.D. (2018). Real-space refinement in PHENIX for cryo-EM and crystallography. Acta Crystallogr D Struct Biol 74, 531–544. 10.1107/S2059798318006551.

58. Emsley, P., Lohkamp, B., Scott, W.G., and Cowtan, K. (2010). Features and development of Coot. Acta Crystallogr D Biol Crystallogr 66, 486–501. 10.1107/S0907444910007493.

59. Krissinel, E., and Henrick, K. (2007). Inference of macromolecular assemblies from crystalline state. J Mol Biol 372, 774–797. 10.1016/j.jmb.2007.05.022.

60. Pettersen, E.F., Goddard, T.D., Huang, C.C., Meng, E.C., Couch, G.S., Croll, T.I., Morris, J.H., and Ferrin, T.E. (2021). UCSF ChimeraX: Structure visualization for researchers, educators, and developers. Protein Sci 30, 70–82. 10.1002/pro.3943.

61. Meng, E.C., Goddard, T.D., Pettersen, E.F., Couch, G.S., Pearson, Z.J., Morris, J.H., and Ferrin, T.E. (2023). UCSF ChimeraX: Tools for structure building and analysis. Protein Sci 32, e4792. 10.1002/pro.4792.

62. Zelter, A., Bonomi, M., Kim, J.O., Umbreit, N.T., Hoopmann, M.R., Johnson, R., Riffle, M., Jaschob, D., MacCoss, M.J., Moritz, R.L., and Davis, T.N. (2015). The molecular architecture of the Dam1 kinetochore complex is defined by cross-linking based structural modelling. Nat Commun 6, 8673. 10.1038/ncomms9673.

63. Chambers, M.C., Maclean, B., Burke, R., Amodei, D., Ruderman, D.L., Neumann, S., Gatto, L., Fischer, B., Pratt, B., Egertson, J., et al. (2012). A cross-plarorm toolkit for mass spectrometry and proteomics. Nat Biotechnol 30, 918–920. 10.1038/nbt.2377.

64. Hoopmann, M.R., Zelter, A., Johnson, R.S., Riffle, M., MacCoss, M. J., Davis, T. N., and Moritz, R. L. (2015). Kojak: efficient analysis of chemically cross-linked protein complexes. Journal of proteome research 14, 2190–2198. 10.1021/pr501321h.

65. Kall, L., Canterbury, J.D., Weston, J., Noble, W.S., and MacCoss, M.J. (2007). Semi-supervised learning for peptide idencficacon from shotgun proteomics datasets. Nat Methods 4, 923–925. 10.1038/nmeth1113.

66. Riffle, M., Jaschob, D., Zelter, A., and Davis, T.N. (2019). Proxl (Protein Cross-Linking Database): A Public Server, QC Tools, and Other Major Updates. Journal of proteome research 18, 759–764. 10.1021/acs.jproteome.8b00726.

67. Meyer, H.J., and Rape, M. (2014). Enhanced protein degradation by branched ubiquitin chains. Cell 157, 910–921. 10.1016/j.cell.2014.03.037.

68. Lazar, G.A., Desjarlais, J.R., and Handel, T.M. (1997). De novo design of the hydrophobic core of ubiquitin. Protein Sci 6, 1167–1178. 10.1002/pro.5560060605.

69. Williamson, M.P. (2013). Using chemical shit perturbation to characterise ligand binding. Prog Nucl Magn Reson Spectrosc 73, 1–16. 10.1016/j.pnmrs.2013.02.001.

